# Meta-analysis of preclinical pharmacogenomic studies to discover robust and translatable biomarkers of drug response

**DOI:** 10.1101/2022.10.22.513279

**Authors:** Petr Smirnov, Sisira Kadambat Nair, Farnoosh Abbas-Aghababazadeh, Nikta Feizi, Ian Smith, Trevor J. Pugh, Benjamin Haibe-Kains

## Abstract

Preclinical pharmacogenomic studies provide an opportunity to discover novel biomarkers for drug response. However, pharamcogenomic studies linking gene expression profiles to drug response do not always agree on the significance or strength of biomarkers. We apply a statistical meta-analysis approach to 7 large independent pharmacogenomic studies, testing for tissue-specific gene expression markers predictive of response among cancer cell lines. We found 4,338 statistically-significant biomarkers across 8 tissue types and 34 drugs. Significant biomarkers were found to be closer than random to drug targets in a gene network built on pathway co-membership (average distance of 2 vs 2.9). However, functional relationships with the drug target did not predict reproducibility across studies. To validate these biomarkers, we utilized 10 clinical datasets, allowing 42/4338 biomarkers to be assessed for clinical translation. Of the 42 candidate biomarkers, the expression of *ODC1* was found to be significantly predictive of Paclitaxel response as a neoadjuvant treatment of breast carcinoma across 2 independent clinical studies of ***>***200 patients each. We expect that as more clinical transcriptomics data matched with response are available, our results can be used to prioritize which genes to evaluate as clinical biomarkers of drug response.

## Introduction

Genome-informed precision medicine programs strive to improve patient outcomes by assigning treatments most likely to elicit a favourable response based on the particular molecular aberrations found in a patient’s tumour(s). While studies have shown benefits to genomically-matched treatment over unmatched therapy [1–3], the rates at which patients can be matched to treatment vary between study design [5-77%], and many studies report finding actionable aberrations in less than half the patients profiled [1, 4–6]. Additionally, a large fraction of anti-cancer therapies lack clear biomarkers predictive of sensitivity or resistance. For example, the OncoKB database of predictive biomarkers includes associations for only 113 therapies as of 2022 [7], out of *>* 200 FDA-approved therapies for oncology [8]. Therefore, many therapies cannot be considered for genomically-matched treatment. Furthermore, while OncoKB identifies 692 cancer associated genes, only 71/692 are associated with response to a cancer therapy. While other databases may vary in the exact numbers, it is clear that we have an incomplete picture of how the molecular characteristics of a tumour affect response to cancer therapy. In turn, this affects our ability to deliver the benefits of genomically-matched therapy to all patients.

*In vitro* pharmacogenomics studies combine drug screening with molecular profiling of cancer models, usually immortalized cancer cell lines, with the goal of discovering new associations between molecular features and drug response. Since 2012, 10 major such studies have been published, discovering novel putative biomarkers of response and made large datasets available for the research community[9–21]. Molecular profiling strategies have varied across theses studies, including mutations, copy number aberrations, transcript expression, methylation state, and (phospho-)proteomics. Even so, these studies have all concluded that gene expression was the most predictive single data modality for the greatest number of drugs [12, 22, 23]. In parallel, recent clinical studies are evaluating the benefit of integrating transcriptomic profiling into the standard precision oncology workflow. Clinical studies of mixed tumour types from the DKFZ, BC Cancer Agency and the WIN Consortium have found that incorporating RNA profiling increases the number of patients successfully matched to therapy [24–26], and patients with RNA-based assignment show similar rates of clinical benefit to those with mutation-based therapy assignment [24, 26]. Furthermore, in studies where DNA evidence was not preferentially used for treatment matching over transcriptomic evidence, transcriptomics has been the most common data modality for finding actionable evidence which can influence treatment decisions [24, 26]. These trials also provide evidence that transcriptomic profiling is feasible in a precision oncology context. Altogether, these results suggest transcriptomics to be a promising modality for discovering novel predictive biomarkers with the potential of increasing matching rates and overall patient benefit in precision oncology trials.

Unfortunately, translation of transcriptomic biomarkers from preclinical prediction to clinical utility has been impeded difficulty reproducing preclinical findings *in vitro* discoveries. An early comparison between two major preclinical pharmacogenomic datasets raised concerns that inconsistent profiling of drug response data lead to inconsistent biomarker discovery across the two studies [27]. This led to scientific debate and efforts to identify sources of inconsistencies, with both different experimental and analytic decisions across studies explaining some of the observed variance [11, 28–30]. While standardizing normalization and analysis approaches, and choosing more appropriate statistical measures, has been able to increase the consistency across studies [31, 32], cross-centre differences in experimental protocols, including different seeding densities, cell viability assays and culture conditions, explain much of the variability among drug response quantification [11, 30]. Such differences across studies are inevitable, as scientific considerations driven by different questions asked of the data, practical considerations driven by available technologies, and the development of novel assays and multiplexing approaches [21] lead to different optimal choices for each study. As such, we believe analytical approaches that are designed to take the resulting variability in biomarker correlations with drug response into account are necessary to discover reproducible and translatable biomarkers from preclinical pharmacogenomics data.

We hypothesized that differences between studies can be turned into an asset, allowing us to identify those biomarkers which are robust to changes in experimental conditions, and therefore, possibly more robust to differences between *in vitro* cell culture and *in situ* tumours. To this end, we apply statistical meta-analysis models to find the biomarkers robust to variation across studies. Here, we present a framework for meta-analysis of preclinical pharmacogenomic studies to discover univariable transcriptomic biomarkers of drug response. We focused on tissue-specific, single gene markers of drug response, which most closely resemble the mutation biomarkers currently widely used in precision oncology trials. Our method evaluates heterogeneity in the estimated effects between datasets, and applies either a fixed- or random-effect linear model to estimate a consensus correlation between expression and drug response. Adapting recent statistics and econometrics literature, we used permutation and bootstrapping approaches to statistical inference to avoid distributional assumptions that may be violated by real world drug-response data. We applyed our framework in a meta-analysis of seven large independent drug screening datasets, creating a list of putative biomarkers. We then conducted a systematic validation of our discoveries in clinical data, building on the recent work by Liu et al. in assembling a resource of clinical datasets with treatment response and transcriptomic information [33], as well as the datasets used for validation of the SELECT method by Lee et al. [34]. We discovered a novel preclinical marker predicting response to paclitaxel as a neoadjuvant therapy among breast cancer patients, demonstrating the potential for translation of our findings. Ultimately, we expect that our framework will establish meta-analysis as a standard step in the analysis of *in vitro* pharmacogenomic data, and that our preclinical discoveries will form a basis for further translational research.

## Results

### Collection of a compendium of preclinical cell line screening datasets

To perform statistical meta-analysis of gene expression-drug associations, we identified seven independent published drug screening datasets with paired independent transcriptomic profiling of cancer cell lines. Three of these seven datasets, the gCSI [11], GRAY [18, 19, 35] and UHNBreast [36, 37] drug screens were unambiguously paired with transcriptomic profiling through RNA sequencing done by the same study authors. Two datasets originated from the Genomics of Drug Sensitivity in Cancer (GDSC) project, corresponding to two iterations of their drug screening platform (GDSC1,GDSC2) [12, 38, 39]. For these screens, two transcriptomic datasets generated by Affymetrix microarray profiling by the GDSC project were paired with drug response data from the GDSC study. The earlier HG-U133A arrays paired to the GDSC1 drug screen, and the HG-U219 array paired with the GDSC2 screen. While RNA sequencing data was also available from the GDSC project, the microarray profiling was preferred due to larger sample sizes. For the CTRPv2 drug screening data, we follow the Cancer Therapeutics Response Portal study authors in pairing their drug screening dataset with RNA sequencing data from the Cancer Cell Line Encyclopedia (referred to from now on as the CCLE.CTRPv2) [10, 16, 40, 41]. Finally, for the drug screening dataset published by the Cancer Cell Line Encyclopedia (CCLE), the data was paired with the original microarray profiles generated by the CCLE group [9]. As such, each of the seven drug screening datasets were paired with independent profiling of expression in the same cells (Supplementary Figure 1).

The drug screening and transcriptomic datasets were downloaded from the ORCESTRA platform [42] and reprocessed with uniform pipelines as described previously [43, 44]. Drug response for each drug-cell pair in each study was quantified using the Area Above the Curve (AAC) metric. Gene expression was analyzed either on a log-RMA normalized scale for microarray, or a log(TPM+0.001) scale. Further filtering of drug screening data was performed to standardize tested concentration ranges within each study (see Methods).

### Implementation of meta-analysis pipeline for univariable transcriptomic biomarker discovery

Having collected seven studies of independent gene expression and drug response profiling across different labs and different collections of cell lines, we set out to discover which gene expression-drug associations were robust and reproducible among studies. While the most commonly used meta-analysis techniques focus on integrating summary statistics (usually effect size and standard error) across studies, having access to the full data from each pharmacogenomics study allowed applying meta-analysis models at the individual sample level. In our study, this was achieved using linear mixed models to estimate associations between gene expression and drug response.

Previous work, both from our group and others, suggested the Pearson Correlation as more appropriate than non-parametric measures of association for the task of biomarker discovery [45, 46]. However, our previous work also suggested that using asymptotically-derived distributions for testing statistical significance for associations between drug response vectors quantified using the AAC measure is likely to lead to an inflation in false positive discoveries. To confirm that this holds when comparing profiles of gene expression across cell lines with drug AAC values, we ran a permutation simulation on gene-drug associations within the CCLE.CTRPv2 dataset. For this limited simulation, we selected two drugs, paclitaxel and lapatinib, and considered only at the breast cancer cell lines within the CCLE, a cancer where both drugs are applied in clinical practice. Paclitaxel, a microtubule-stabilizing chemotherapy, displays a broad range response across cell lines, while lapatinib, a targeted dual kinase inhibitor of HER2 and EGFR, has an exponential-like distribution where only a few cell lines respond strongly to the drug (Supplementary Figure 2 (a),(b)). To compute the effect size and significance of random associations between gene expression and drug response via Pearson correlation, we then subsampled 1,000 random protein-coding genes from the full gene expression profiles and randomized the drug response data using 10,000 random permutations of the AAC values for the two exemplary drugs. For each calculated correlation, we applied the standard Pearson t-test to reject the null hypothesis of no correlation - which by construction we expect to hold true as the data were permuted. Under the null hypothesis, a test with proper Type I error control is expected to produce p-values uniformly distributed between 0 and 1. However, we observed an inflation of p values towards smaller values, which was more pronounced for Lapatinib than Paclitaxel (Supplementary Figure 2 (c),(d)). Visually examining the results of the permutation test for specific genes, we notice that for both drugs there are examples of genes for which the correlations with permuted labels had an inflation of low p values (Supplementary Figure 2 (e),(f)). Therefore, we concluded that for any particular association between gene expression and drug response, the Type I error control would vary depending on the particular distribution of expression and drug response observed. To avoid the demonstrated inflation in Type I errors associated with the standard t-test, we adopted resampling methods, such as permutation and bootstrap testing, for robustly evaluating the significance of associations.

One major drawback of resampling methods for evaluating statistical significance is the computational cost. This is especially true when testing hypotheses genome-wide, as the p value cutoff (*α*) after multiple testing correction can be in the range of 10^−6^ − 10^−8^. Estimating p values using resampling methods to such small precision can require millions to billions of samplings from the observed data for each hypothesis tested. We therefore applied two strategies to alleviate this computational cost and make a transcriptome-wide analysis feasible. The first was the use of a published adaptive permutation testing approach termed QUICK-STOP [47], which stops a permutation test as soon as enough evidence is observed to decide on the significance of a particular test with a predetermined level of confidence. The second strategy was to focus our meta-analysis only on those markers for which a significant association was found within at least a one study. As our study was primarily focused on evaluating single-expression feature markers of response, our focus was on finding the strongest biomarkers for each drug-tissue combination, and not necessarily on an exhaustive enumeration of features associated with drug response. Therefore, while meta-analysis can be employed to combine several studies displaying weak evidence for an association, for the purpose of our study, meta-analysis was used primarily to ensure robustness across experimental conditions for the associations discovered. Permutation testing associations within each study was significantly quicker than testing the meta-analysis through permutation or bootstrap techniques, and therefore applying a filter requiring all markers examined in meta-analysis to have shown a significant correlation with drug response within at least a single dataset provided an efficient method to reduce the computation necessary to complete this analysis.

Our pipeline for evaluating univariable expression markers therefore consisted of three stages. For a given drug and tissue of origin, the first stage consists of adaptive permutation testing the significance of the Pearson correlation between the expression of each gene and drug response (measured in AAC) within each study separately. Each gene expression-tissue-drug triplet significant in at least one study then underwent the next two meta-analysis stages. In the second stage, the permutation test method of Lee and Braun [48] was used to detect heterogeneity of effects across studies for each gene-tissue-drug triplet. Gene-tissue-drug triplets which had a heterogeneity test p-value of greater than 0.1 were deemed to not show evidence of random variation between studies, and in the third stage were tested with a fixed effect model, which fits a common correlation coefficient between gene expression and drug response across all studies. Gene-tissue-drug triplets which had a heterogeneity test p-value of less than 0.1 were tested in the third stage using a random effects model, which allows for random variation of effects across studies. In this case, inference was done on the mean of the estimated correlation distribution across studies. Permutation and bootstrap tests were used to evaluate the significance of each association for the fixed and random effect models respectively. In both cases, the effect size was measured using Pearson correlation (details in methods). Multiple testing correction was applied using Bonferroni’s method, accounting for the total number of gene-expression features examined in the first stage. We deemed associations that were significant after multiple testing correction to be putative biomarkers for further investigation.

In applying this pipeline to the seven datasets described above, we required a drug to have been tested on at least 20 cell lines within a particular tissue type across at least three studies for that drug-tissue combination to be considered. This filter left us with 70 compounds tested across a subset of 11 tissue types, forming 483 drug-tissue combinations for our study. We then similarly filtered to genes whose expression was measured in at least 20 cell lines within each tissue type under consideration. This left us with a total of 22,778,865 tissue-compound-gene associations to consider in our analysis.

### Meta-analysis of tissue specific associations reveals robust biomarkers of drug response

We then applied our pipeline to the 22,778,865 candidate gene-drug-tissue associations that could be evaluated in at least three datasets. We found 7,804 associations between gene expression and drug response significant in at least one dataset (Supplementary Table 1). Of these, approximately 56% (4,338) across all datasets remain significant after meta-analysis (Supplementary Table 1). Significant biomarkers were found in 8 different tissue types, predicting response for 34 different drugs. This was fairly consistent regardless of which dataset the association was detected in, with between 38% and 66% of discoveries in each particular dataset not validating in meta-analysis (Figure 1b). This suggests that if any individual dataset was used for biomarker discovery in isolation, nearly half the discoveries would not reproduce across validation studies.

**Fig. 1:**
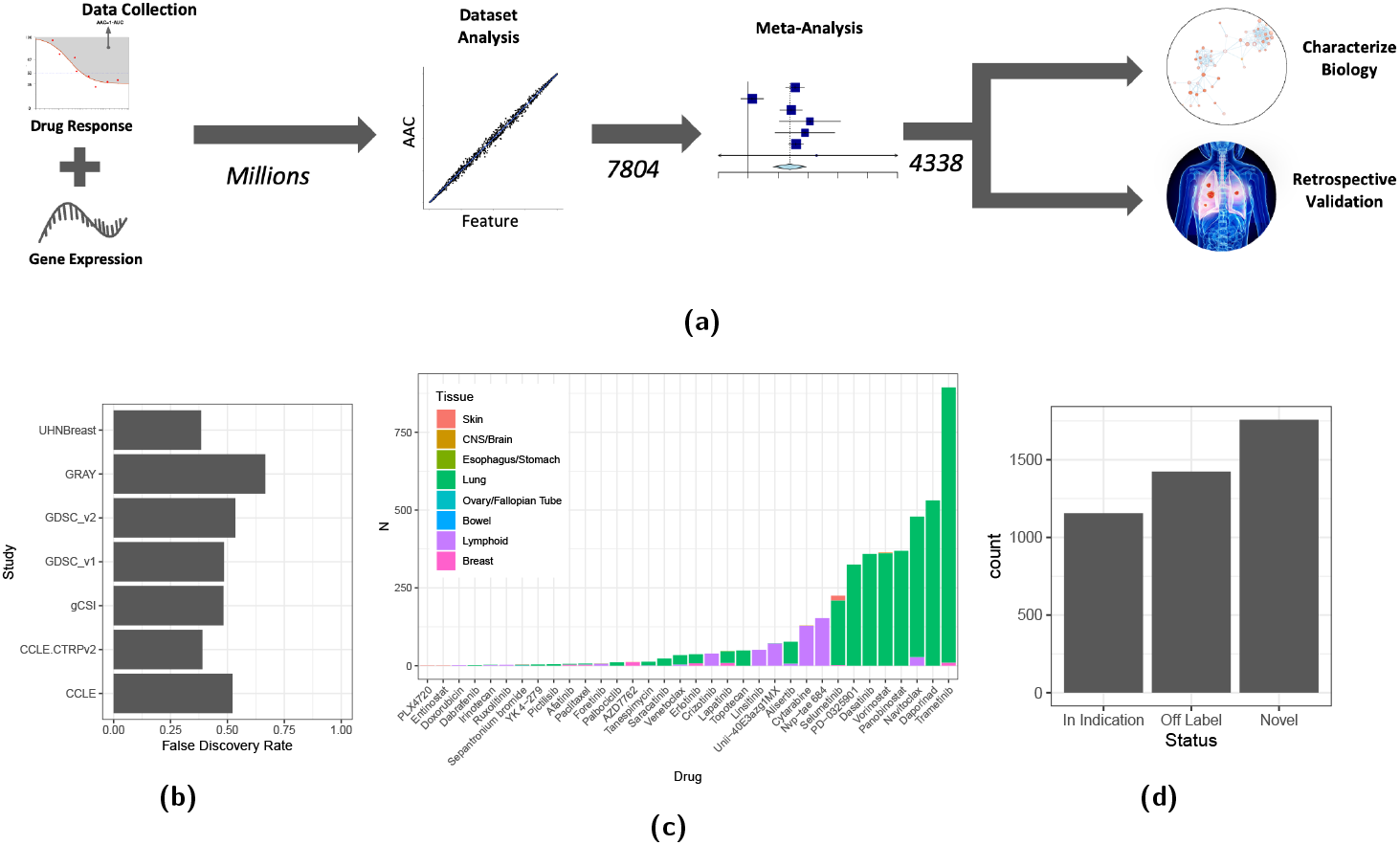
**a)** A visual diagram of the meta-analysis pipeline and study design; **b)** The rate at which discoveries made in each individual dataset fail to be confirmed as significant associations after meta-analysis; **c)** the number of biomarkers found for each drug, coloured by the tissue type of the cell lines among which each biomarker was discovered; **d)** the distribution of biomarkers found per the status of the drug-indication for which they were discovered. In Indication = drug is currently used to treat cancers of this tissue type; Off Label = drug is currently FDA approved for treatment of a cancer, but not used in the tissue type where the biomarker was discovered; Novel = a biomarker for a drug which is not yet used for cancer treatment.

The associations found to be significant after meta-analysis were not uniformly distributed across tissue and compounds (Figure 1c). Rather, there was a pronounced bias towards discovering more associations among lung cancer cell lines. We hypothesized that this can be explained by the differences in sample size across the different tissue types among the seven datasets used for discovery. Larger samples sizes lead to more power to detect significant associations with a smaller effect size, even using resampling based statistical inference. As expected, we find that the minimum absolute value of the significant Pearson correlations across all biomarkers detected is correlated with the number of cell lines in our largest dataset for that tissue type (Supplementary Figure 3a). However, this correlation does not exist for the maximum absolute Pearson correlation (Supplementary Figure 3b). This suggests that the differences in the number of discovered biomarkers between tissues are indeed caused by larger sample sizes allowing us to discover more, weaker associations.

The available data allowed us to discover biomarkers for drugs which are already regularly used in the clinic. 27% of the significant associations after meta-analysis were with FDA approved compounds and in a tissue type encompassing at least one cancer for which the compound is currently indicated, according to the FDA approved drug labels (Figure 1d). A further 33% of the associations were found for FDA compounds in tissue types where they are not indicated. The first set of biomarkers can in principle be investigated directly in the clinic today, by collecting RNA expression data with patients already being treated with these drugs. The second set can suggest subgroups of cancers in tissues where compounds are not indicated, but which may benefit for an off label application of the compound, and therefore may still be relevant in the context of precision oncology trials.

### Known cancer-related genesets are enriched for biomarkers, but are not predictive of consistency across datasets

Genome-wide expression microarrays and RNA sequencing have allowed the unbiased profiling of transcript expression, without the need to pre-select particular genes or transcripts of interest. A common approach to reduce multiple testing burden and increase statistical power in analyses of such data has been to use independent criteria to filter down the explored feature space to a more manageable number of expression features prior to hypothesis testing. This smaller feature space is chosen to be enriched for features likely relevant for drug response based on *a priori* information - for example, in the case of predicting cancer drug response, genes harbouring known cancer drivers have been used [12].

We were interested in evaluating whether such cancer-related genesets would be enriched for significant associations. To answer this question, we used two cancer-related gene lists: The Cancer Gene Census from COSMIC [49], as well as the list of genes harbouring Cancer Functional Events [12]. We note that these genesets were originally defined using genetic and epigenetic evidence of association with cancer, not based on their expression. We also include in our analysis the union of all genes defined in the Hallmark genesets from MSigDB [50–52]. While not strictly cancer related, this set of genes is comprised of members of well-characterized biological processes and would be a sensible feature space to enrich for easily interpretable results.

Comparing the results of our meta-analysis to the examined genesets, we found that all three are enriched in biomarkers (Supplementary Figure 4a). While the observed odds ratios were all significantly different from 1, as well as from each other, the enrichment was remarkably higher for the Cancer Gene Census genes, with an observed OR=8.94 vs 3.10 and 2.51 for the Hallmark and Cancer Functional Events respectively. Overall, these results confirm that filtering to features *a priori* associated with cancer can enrich for biomarkers of drug response while reducing multiple-testing burden. However, such filters may not always be appropriate, as they limit the ability to discover biomarkers among genes not previously associated with cancer.

We further investigated whether the same genesets have any relationship with the reproducibility of a biomarker across different *in vitro* studies. We tested whether these genesets are enriched in markers that validate in meta-analysis, among those markers that were significant in at least 1 individual dataset. Interestingly, we found no sign of enrichment for biomarker robustness among the Cancer Gene Census genes (OR=1.06, p=0.5), and only weak trends towards enrichment for the Hallmarks (OR=1.1, p=0.05) and Cancer Functional Events (OR=1.20, p=0.08) (Supplementary Figure 4b). Although cancer-related genesets are enriched in genes associated with drug response, there is no evidence that these genesets select for associations robust across experimental conditions and datasets. Therefore, while analysis can be simplified and statistical power increased by pre-selecting cancer-related or well-characterized genesets, meta-analysis is still a necessary step to ensure robustness and reproducibility of the discovered associations.

### Functional proximity to drug targets is not a predictor of biomarker strength

We next evaluated whether expression biomarkers of drug response were enriched for genes that had a functional relationship with the annotated drug targets. Of the 34 drugs for which at least one significant biomarker was found after meta-analysis, 32 had at least one annotated drug target. Using a gene-gene network based on co-membership in Reactome pathways, we calculated for each drug the minimum distance in this network between each biomarker and any annotated drug target. We then compared the distribution of these distances for each biomarker to two backgrounds: the distribution of distances between any random pair of genes in the network, and the distribution between a randomly selected gene and a drug target (Figure 2a). We found that the average distance of a random gene to a drug target is significantly smaller (Wilcoxon rank-sum p value: 3.02 · 10^−13^) than the distance between two random genes, suggesting that drug targets are enriched for genes central in this functional network. Biomarkers of drug response were found to be closer still on average to drug targets (Wilcoxon rank-sum p value: 3.37· 10^−16^). However, the majority of biomarkers remained at least a distance of 2 away in this network (compared to an average distance between two genes of 2.9). There was also a significant association between a biomarker’s absolute Pearson correlation and its distance within the network (Kruskal-Wallis p value: 2.45· 10^−5^), however, we found meaningful differences in the association strength only at the two extremes (Figure 2b). For biomarkers of drug response which themselves were an annotated drug target, there was a significant increase in the average biomarker strength compared to all other biomarkers (Wilcoxon rank-sum p value: 0.044). On the other end, markers which were a distance of 5 edges away, the largest observed distance, were weaker on average, although the difference was not significant. We did not, however, observe any remarkable differences between biomarkers which shared pathway co-membership with drug targets (distance of 1) compared to biomarkers which were a distance of 4 away.

**Fig. 2:**
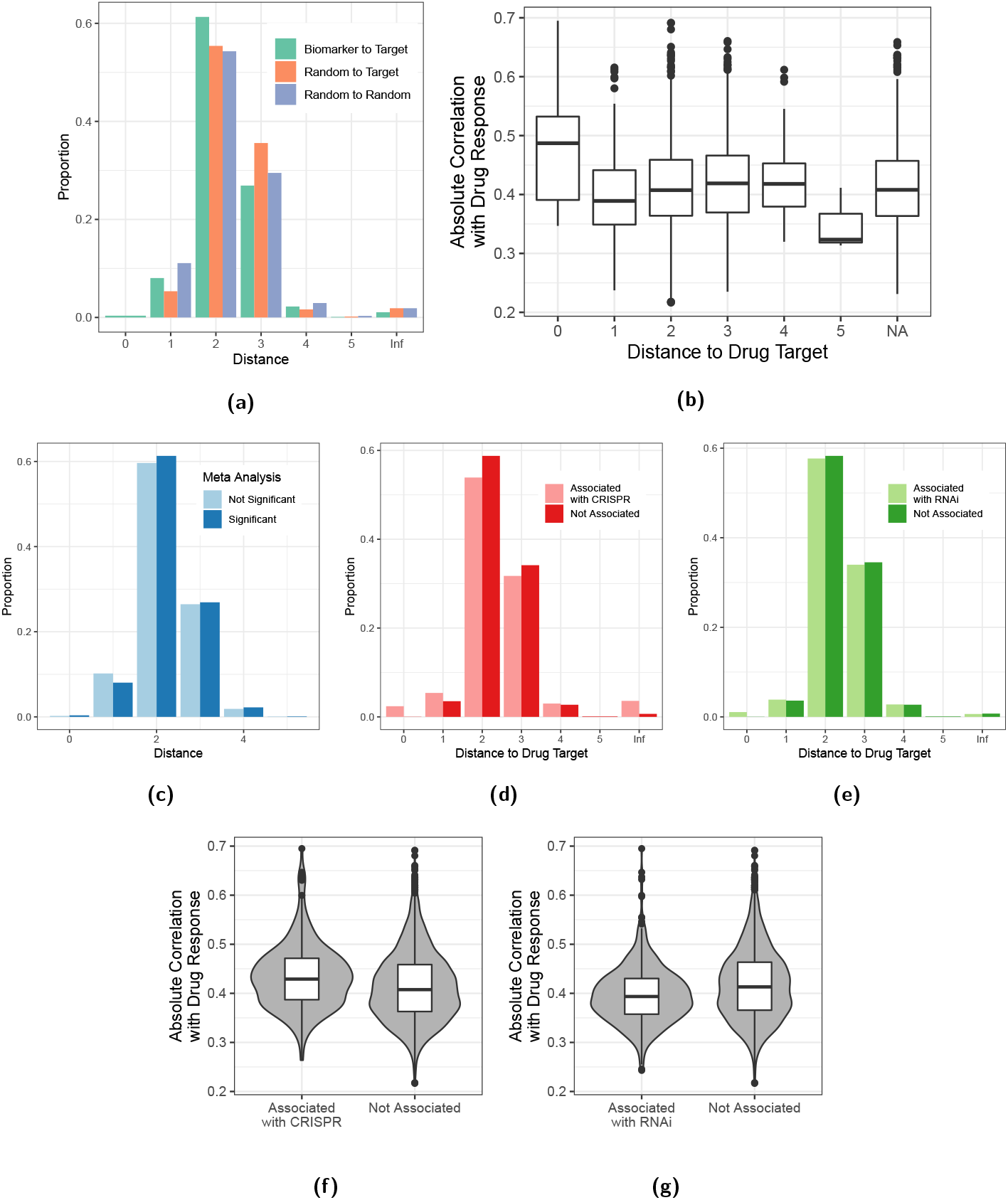
**a)** Distribution of distances in the Reactome pathway-based gene-gene network from biomarkers to drug targets, all genes to drug targets, and between pairs of random genes. Infinite distances indicate that there is no path in the network between the biomarkers and drug targets; **b)** Biomarker effect size as a function of distance to the drug target in the Reactome network. While the effect of distance on the mean effect size is significant (Kruskal Wallis p: 2.45 · 10^−5^), only biomarkers at a distance of 0 (cases where the drug target is the biomarker) show a significant difference from the rest (Wilcoxon rank-sum test p: 0.044); **c)** distribution of distances in the Reactome network between dataset-level biomarkers and drug targets of the respective drug split by whether a biomarker passes or fails meta-analysis; **d), e)** Distribution of the distance to each annotated drug target for biomarkers of the respective drug, split by whether the biomarker is also correlated with CRISPR and RNAi inhibition of the same drug target respectively; **f), g)** Absolute correlation of biomarker with drug response for biomarkers with and without association to CRISPR and RNAi respectively.

While proximity to drug targets enriched for biomarkers among random genes, it failed to predict whether a biomarker discovered in a dataset would validate after meta-analysis. Comparing the distribution of distances to a drug target for biomarkers significant after meta-analysis to those genes which were significant in 1 dataset but not in meta-analysis, we found no significant differences (Wilcoxon rank-sum p value: 0.24). Taken altogether, our results suggest that drug response is a complex phenotype that is not completely predicted by biological processes directly related to the annotated drug targets. This may occur due to the effect of distantly related biological processes on the ultimate cell fate, underappreciated polypharmacology of the compounds in our study, or incomplete understanding and annotation of biological pathways, especially in the context of cancer, supporting the need for unbiased search when looking for expression markers or signatures of drug response.

Inspired by the idea of “robust biomarkers” as defined by Gonçalves et al. [53], we next evaluated whether correlation with annotated drug target gene inhibition through CRISPR or RNAi would enrich for gene expression biomarkers using the Cancer Dependency Map datasets [15, 54–58]. For each RNAi or CRISPR inhibition of an annotated drug target, we computed tissue-specific correlations between biomarkers of response for associated drugs and the essentiality scores. We observed that only 15% of biomarkers also predicted response to RNAi of the drug targets, and 6% predicted CRISPR knockdown response. We compared the Reactome pathway graph distance between biomarkers which correlated with inhibition of a particular drug target and those without a significant correlation to CRISPR or RNAi of that target. While there was a trend towards a shorter average distance for biomarkers associated with both CRISPR and RNAi (Figures 2d, 2e), neither trend was significant (Wilcoxon rank-sum p values: 0.83 and 0.53 respectively). We found significant but small differences between the strength of these two groups of biomarkers (Figure 2e, 2f), but the direction disagreed between RNAi and CRISPR associated markers (Wilcoxon rank-sum p value: 1.64·10^−12^ and 1.74·10^−6^ for RNAi and CRISPR respectively). Significant correlation of a dataset-level biomarker with RNAi or CRISPR inhibition also failed to enrich for biomarkers which validated after meta-analysis. We found that genes correlating with RNAi were neither more nor less likely to pass meta-analysis (Odds Ratio: 1.05, Fisher test p value: 0.49), while correlation with CRISPR-based drug-target inhibition actually moderately enriched for genes which did not pass meta-analysis (Odds Ratio: 0.61, Fisher test p value: 2.18·10^−08^). Overall, our results suggest that predicting RNAi and CRISPR inhibition of drug targets may enrich for biomarkers that can be interpreted in the context of known drug mechanism of action, but it does not have a meaningful difference on the predictive strength of the biomarkers found *in vitro*, or in their reproducibility across different studies.

### *In vitro* markers of drug response among lung cancer cell lines recapitulate known vulnerabilities of neuroendocrine-derived lung cancers

As already observed above, the majority of significant associations were found after meta-analysis among lung cancer cell lines. While significant associations were found with 24 drugs with 1574 unique genes, these were not equally distributed among different compounds. While the median number of biomarkers per drug was 29.5 in the lung tissue type, for eight of the drugs we found ≥ 100 significant biomarkers of response after meta-analysis. Interestingly, many of the genes predicting response to these compounds were common across multiple compounds, suggesting a global expression pattern may be driving differential response to drugs across lung cancer cell lines. We therefore selected for those markers that were associated with at least four of these eight drugs, and clustered the drugs using hierarchical clustering on a cosine distance between their biomarker signatures. We observed that the eight drugs cluster tightly into two groups of four by their shared markers of drug response (Figure 3a). This clustering was replicated when clustering all 24 drugs across all 1574 genes, with most other drugs clustering into degenerate clusters, as they shared no or few biomarkers with each other (Supplementary Figure 5). The exceptions were erlotinib, afatinib and lapatinib forming a HER-family inhibitor cluster, irinotecan and topotecan clustering together, and palbociclib clustering with tanespimycin.

**Fig. 3:**
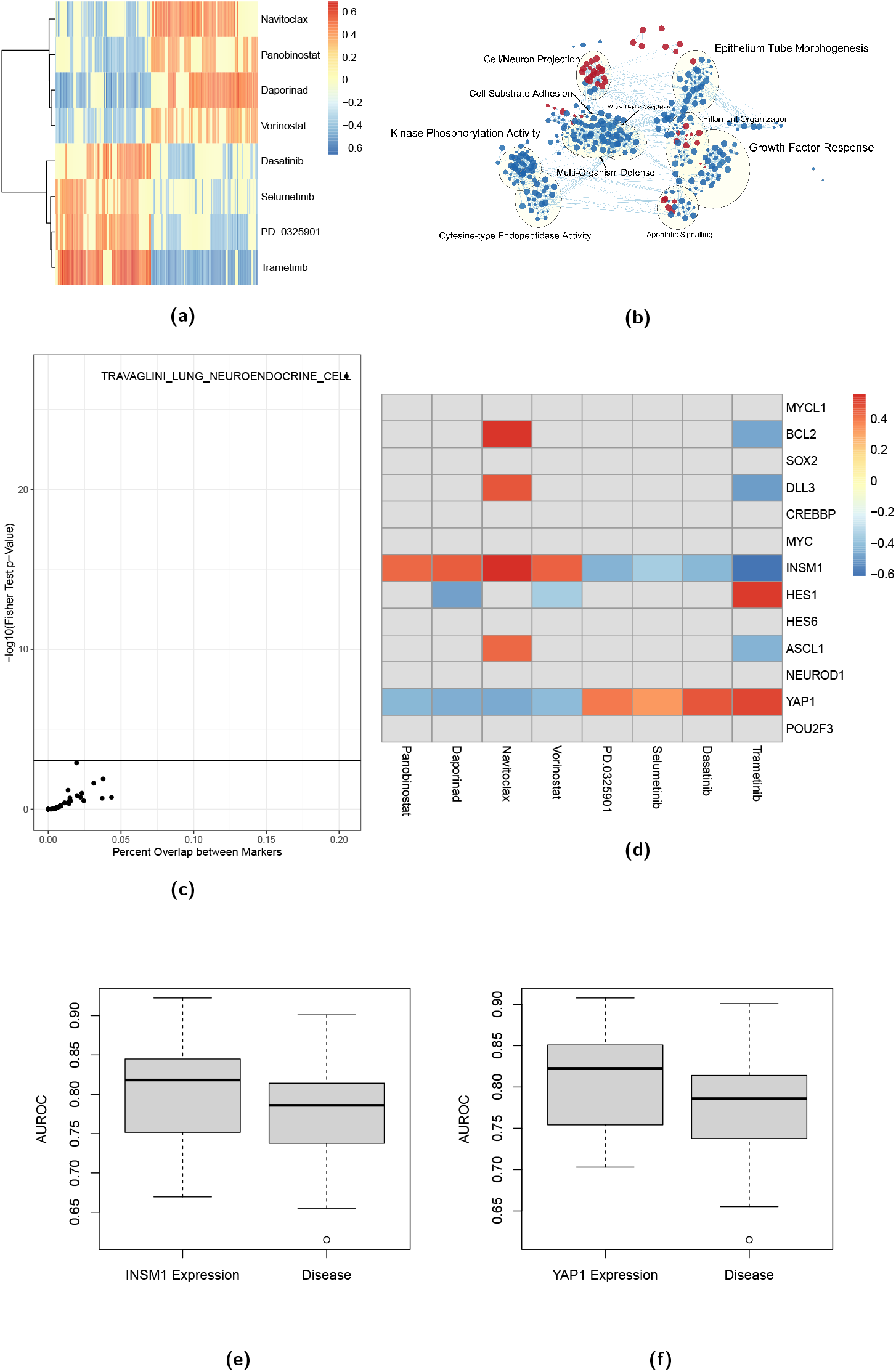
**a)** Heatmap of the associations between gene expression (rows) and drugs (columns) after meta-analysis among lung cancer cell lines, for those drugs with ≥ 100 biomarkers and those biomarkers predictive of ≥ 4 drugs. Values of 0 indicate a that a gene was not significantly associated with drug response after meta-analysis. Wald D2 clustering on correlation distance is displayed for the rows; **a)** An Enrichment Map of over-represented GO Biological Process terms among biomarkers of sensitivity (red) or resistance (blue) to the top cluster (cluster 1) of **a)**; **c)** A volcano plot of the overrepresented lung cell-type markers as defined by Travaglini et al. among biomakers predicting sensitivity to cluster 1 drugs; **e)** A heatmap of associations between known marker genes of Small Cell Lung Cancer (SCLC) subtypes and response to the 8 drugs in **a)**; **e), f)** Comparison of the AUROC comparing continuous drug response (AAC) to either binarized high/low expression of INSM1 or YAP1, or Disease labels (Non-SCLC vs SCLC), among the lung cancer cell lines across the CCLE.CTRPv2, GDSC1 and GDSC2 datasets and 8 drugs from **a)**.

We were interested in further characterizing the two clusters of drugs driven by ≥ 100 biomarkers. The first drug cluster consisted of navitoclax, panobinostat, vorinostat and daporinad, a BCL2 inhibitor, two HDAC inhibitors and a NMPRTase inhibitor. The second cluster consisted of dasatinib, selumentinib, trametinib and PD-0325901, a BCR/ABL inhibitor and three MEK inhibitors. Of 324 genes significantly associated with response to at least four of these drugs, only 8 were not also associated with opposite responses (sensitivity/resistance) to at least one drug in each of the two clusters. Therefore, we considered the intersection of genes predicting sensitivity to cluster 1 and resistance to cluster 2 drugs as the Cluster 1 marker geneset, and the intersection of genes predicting sensitivity to cluster 2 and resistance to cluster 1 drugs as the Cluster 2 marker geneset.

To identify which biological processes were involved in driving this difference in drug response phenotypes, we performed over-representation analysis for GO Biological Processes among the Cluster 1 and Cluster 2 marker genesets (Supplementary Table 2). 71 GO terms were over-represented among the genes associated with Cluster 1 sensitivity, and 424 among genes associated with Cluster 2 sensitivity. We then used Enrichment Map to visualize the relationships between enriched GO terms, and applied network clustering to identify clusters of enriched GO terms (Figure 3b). GO terms related to Kinase Phosphorylation Activity and Growth Factor Response were enriched among markers of resistance to cluster 1/sensitivity to cluster 2. Interestingly, Cell Substrate Adhesion terms were also associated with resistance to cluster 1 drugs, and at the same time terms associated with Cell/Neuronal projection were enriched among markers of cluster 1 drug sensitivity. Small Cell Lung Cancer cell lines commonly show a non-substrate adhesive preference for growth [59], and many Small Cell Lung Cancers are thought to originate from a neuroendocrine cell of origin, suggesting that the difference in drug response observed may be partially explained by different response among cancers with a different cell of origin within the same tissue.

We repeated the over-representation analysis using markers of cell types identified from the single cell profiling of lung tissue by Travaglini et al [60]. While markers of resistance to cluster 1/sensitivity to cluster 2 were enriched in markers of multiple cell types including epithelial, but also immune cells, markers of sensitivity to cluster 1/resistance to cluster 2 were enriched strongly in markers of neuroendocrine cells (Figure 3c and Supplementary Figure 6). Neuroendocrine tumours of the lung are accepted to be Small Cell Lung Cancers (SCLC) in histology, but not all SCLC tumours express markers of neuroendocrine lung cells. Of the four accepted subtypes of Small Cell Lung Cancer, two are considered to be neuroendocrine-like, and are marked by high expression of *INSM1* and low expression of *YAP1* [59]. We therefore examined whether these subtype markers, as well as other genes known to be associated with subtypes of SCLC were among the markers identified in our meta-analysis. *INSM1* was found to be among the markers predicting response to cluster 1 drugs/resistance to cluster 2 drugs, with *YAP1* showing the opposite association (Figure 3d). Expression of ASCL1, DLL3 and BCL2, markers of the first subtype of SCLC were found to be associated with Navitoclax response and Trametinib resistance. We then went back and examined the relationship between expression of *INSM1* and *YAP1* with drug response for these eight drugs across the three largest datasets in our study (Supplementary Figure 7,8) stratified by cell lines labeled as originating from non-small cell and small cell lung cancers separately. As expected, *INSM1* and *YAP1* expression was observed to be strongly correlated with NSCLC/SCLC labels. However, cells from either disease that expressed these markers more similarly to the average of the other disease also tended to display response more characteristic to the other disease type. We therefore hypothesized that *YAP1* and *INSM1* expression would be a more correlated with response to the eight compounds than disease of origin labels for the cell lines.

To compare between the continuous *YAP1* /*INSM1* expression and disease labels, we assigned to each cell line a *YAP1* high/low and *INSM1* high/low expression label based on a Mixture of Gaussians model of gene expression. We then compared the AUROC between AAC and binary expression or disease labels. For both *YAP1* and *INSM1*, we found that AAC was a better predictor of expression classes than disease type labels for 18/22 drug-dataset pairs (Figure 3e,3f and Supplementary Figures 9,10). Taken together, these results suggest that differences in the response to drugs between the two identified clusters are driven by a neuroendocrine cell phenotype, and expression of neuroendocrine markers is more predictive than disease classification for response to these compounds. This may be clinically relevant as both HDAC inhibitors and BCL2 targeted drugs are currently under investigation as novel treatments for Small Cell Lung Cancers [61–64], with existing preclinical evidence suggesting that the sensitivity is specific to neuroendocrine subtypes of SCLC [59, 62].

### Identification of clinical datasets for retrospective validation of predictive biomarkers

While preclinical pharmacogenomics experiments can be used to characterize the molecular basis of response to therapies, translation of these findings into clinical predictors will require evidence from *in vivo* model systems or retrospective clinical data. As such, we conducted a search for published clinical datasets combining treatment response/post treatment survival and gene expression profiling. For this search, we leveraged two recent resources: the CTR-DB database [33], and the datasets used by Lee et al. [34] for validation of the SELECT method (Supplementary Table 3). For each biomarker, we selected the clinical datasets which matched both tissue type and drug, either in monotherapy or combination. Unsurprisingly, retrospective clinical data were only rarely available for our biomarkers (Supplementary Figure 11), as only 5 drug-tissue combinations had retrospective data available: Paclitaxel and Lapatinib treatments in Breast Cancer, Erlotinib treatment in Lung Cancer (NSCLC), Paclitaxel treatment in Ovarian Cancer and Doxorubicin treatment in cancer of Lymphoid cells (T-ALL).

### Filtering of biomarkers unlikely to be conserved between cell lines and tissues

Given the many differences between immortalized cancer cell lines and *in situ* tumours, it is expected that the expression of genes and their relationship with drug sensitivity phenotypes will not be conserved between the two contexts, even for the same cancer types. Published methods exist to “align” cell line and tumour samples [65], or alternatively to find a transformation of gene expression measurements that captures common variation between cell line and tumour expression patterns [66, 67]. However, none of these methods can identify which particular genes are conserved in their expression patterns, or give a probability to predictive single-gene marker translating to the clinical context. To address this gap, we applied a purely statistical method, comparing the distribution of expression for each gene between cell lines and TCGA tumours originating from the same tissue type. We apply the Anderson-Darling test to detect significant shifts in distribution [68], and mark those biomarkers for which the False Discovery Rate is less than 0.1. We compute these statistics genome-wide, for each of the Lung, Ovarian and Breast tissue types where we have clinical validation data.

We find that 47.1%, 27.5%, and 41.1% of genes were detected as having a significant distribution shift by this method for the Lung, Ovarian and Breast tissue types, respectively (Supplementary Table 4). We hypothesize that markers with a significant shift detected by this test are unlikely to validate in clinical data, and should be filtered out prior to validation. Unfortunately, we are unable to test this hypothesis due to our small number of clinical validation datasets only allowing us to examine 33 biomarkers in retrospective clinical data. Nevertheless, we report below whether the biomarkers we are able to examine in retrospective clinical data pass this filter step. We also note that these percentages may be confounded by sample size, and cannot necessarily be interpreted as a measure of how well the cell lines recapitulate tumour biology across different tissue types. However, as we were also able to detect more biomarkers in tissue types with more samples, we find a stricter filter for these tissues appropriate.

### Biomarkers for doxorubicin in lymphoid cell lines and paclitaxel in ovarian cell lines fail to validate in clinical studies

Two biomarkers for doxorubicin response among Lymphoid cells were identified during our *in vitro* meta-analysis: expression of *YIF1A* and *GABRG2* both correlated with resistance to doxorubicin treatment. There was one study in the collection examined where clinical response following doxorubicin treatment was recorded for a cancer of Lymphoid cell types: the study by Winter et al. of transcriptomic predictors of induction failure in T-ALL [69]. This study profiled two independent patient populations using slightly different Affymetrix microarray platforms (50 patients using HGU133Plus2 from the COG 9404 trial and 42 patients using HGU133A from the COG 8704) to compare gene expression profiles between three groups of patients: those that had Complete Continuous Response, those that Relapsed, and those that experienced Induction Failure. These patients were treated with induction chemotherapy consisting of a cocktail of 5 drugs: doxorubicin, mercaptopurine, methotrexate, prednisone and vincristine. We first examined whether *YIF1A* or *GABRG2* expression was predictive of these different categories of response in either study independently. *YIF1A* trended towards a significant association (Kruskal Wallis Bonferroni corrected p: 0.10) with response among the COG 9404 cohort, which was driven by lower expression of *YIF1A* among patients which failed induction other association was significant in either cohort (Supplementary Figure 12a,b,c,d). The direction of the trend among COG 9404 was surprising, as *in vitro*, expression of *YIF1A* predicted resistance, whereas in this cohort, expression correlated with induction of remission by chemotherapy. This association was not reproduced in COG 8704, however only 1 patient in this cohort failed induction. To resolve this inconsistency and increase statistical power, we also analyzed the two cohorts together, pooling the gene expression data after correcting for batch effects between the two chips using ComBat. In this case, neither biomarker was significantly predictive of response (Supplementary Figure 12e,f). The failure of both *in vitro* markers to predict response can be explained in part by the large number of concurrent therapies used in addition to Doxorubicin, as well as the fact that T-ALL derived cell lines were outnumbered by plasma cell leukemia-derived cell lines in the meta-analysis (Supplementary Table 7).

Our *in vitro* analysis revealed a single biomarker of paclitaxel response among ovarian cell lines: higher expression of the *GJB1* gene predicted resistance to paclitaxel. We identified four public datasets with cohorts of ovarian cancer patients treated with either paclitaxel or an unspecified taxane class chemotherapy [70–73]. Out of these four datasets, three had a measure of response for patients, while one dataset had only Overall Survival information for the patients. *GJB1* did not pass the Anderson-Darling filter, and we find no significant association between *GJB1* expression and response among the 3 clinical datasets with response annotation (Supplementary Figure 13a,b,c). For the fourth dataset, we found a trend towards the opposite association, with higher expression of *GJB1* predicting longer survival (Supplementary Figure 13d). Given that *GJB1* codes for a membrane junction protein involved in cell-cell transmission of ions and compounds, its function may be not conserved between *in situ* tumour microenvironments and the cancer cell lines growing in vitro.

### Biomarkers for erlotinib in lung cell lines and lapatinib in breast cancer cell lines trend towards significance in predicting clinical response

Our *in vitro* analysis suggested 29 biomarkers for erlotinib response among lung cancer cell lines. The single available dataset for lung cancer patients treated with Erlotinib comes from the BATTLE1 trial [74], where patients with stage IIIB or higher NSCLC who have failed front line therapy are enrolled into the trial for molecular profiling and possible treatment with targeted therapy matched to the aberrations found. Data were available for 25 patients treated with Erlotinib, with Progression Free Survival (PFS) times ([0.23-5.62] months, median PFS: 1.91 months) and a 100% event rate. We tested the association between expression of the 29 putative biomarkers and progression free survival time using the Concordance Index statistic to measure correlation. Unfortunately, the small sample size meant that none of the biomarkers were significantly associated with response among patients (Supplementary Table 5). Of note, *EGFR* expression, the primary drug target or erlotinib, while being a biomarker *in vitro*, had no association with PFS among patients within the BATTLE1 trial (CI=0.49). When we apply the Anderson-Darling filter for distribution shift, comparing the distribution of the 29 genes between cell lines and TCGA Lung Adenocarcinoma (LUAD) tumours, we find that only 6/29 of these genes have no significant differences in expression detected. If we repeat the validation analysis after filtering to these six markers, we find that expression of one of these six genes, *MAD2L2* is trending towards an association with shorter survival after erlotinib treatment (CI = 0.32, permutation Bonferroni-corrected p value: 0.062) (Supplementary Figure 14a). We further examined whether *MAD2L2* had any purely prognostic value among the TCGA LUAD tumours, and found no significant association with overall survival in this cohort (Supplementary Figure 14b). These results suggest that *MAD2L2* may be a weak marker of erlotinib resistance among NSCLC patients, however, larger sample sizes will be required to make a definitive assessment.

Nine biomarkers for lapatinib response among breast cancer cell lines were identified as significant by our meta-analysis. There was a single dataset available to validate predictive biomarkers for Lapatinib response among Breast Carcinoma (BRCA) patients. The CHER-LOB study evaluated the effectiveness of trastuzumab and lapatinib with standard chemotherapy in comparison to chemotherapy with either arm alone in HER2+ BRCA patients [75]. Pre-treatment biopsies were collected for biomarker analyses, including gene expression profiling using an Affymetrix HGU133 Plus 2 array. This trial contained three arms, all including paclitaxel and FEC (fluorouracil, epirubicin, and cyclophosphamide) chemotherapy in addition to: lapatinib (n=31), trastuzumab (n=23) or trastuzumab + lapatinib (n=34). We evaluated whether any of the *in vitro* biomarkers would predict response in either lapatinib arm separately. Eight out of the nine *in vitro* biomarkers predictive of response to Lapatinib in cell lines were measured on the array used in the clinical dataset. Only expression of *GRB7* trended towards being significantly predictive, and only among the Chemotherapy + lapatinib arm (Wilcoxon rank-sum Bonferroni corrected p value: 0.093) (Supplementary Table 6, Supplementary Figure 15a). Applying the Anderson-Darling filter, comparing the distribution of these eight genes between cell lines and TCGA BRCA tumours, we retain 4/8 *in vitro* biomarkers. If we restricted our analysis to only these genes, then *GRB7* expression would be a significant predictor of sensitivity within the chemotherapy + lapatinib arm. The pCR rate within this arm of the trial was 26%, and *GRB7* expression showed an AUROC of 0.8 and AUPR of 0.57 as a predictor of pCR (Supplementary Figure 15b,c). Overall, there results suggest that GRB7 may be a clinically predictive marker of Lapatinib response in a neoadjuvant setting, however validation in further clinical datasets with larger sample sizes is still required.

### *In vitro* analysis reveals ODC1 as a single gene predictor of neoadjuvant Taxol response among BRCA patients

We found expression of three genes to be predictive of Paclitaxel sensitivity among breast cancer cell lines: *ODC1, EIF5A* and *EIF4A1*. We examined the two largest cohorts of breast cancer patients treated with Paclitaxel for which gene expression profiles of pre-treatment biopsies were available [76, 77]. The first cohort, from the MAQCII study, consisted of 278 patients without any selection by receptor status, and all patients received six months of Paclitaxel plus FAC neoadjuvant therapy, with pathological Complete Response (pCR) assessed at time of surgery. The second cohort, from the Hatzis et al. study, consisted of 478 HER2-BRCA patients, who all received sequential Taxane (Paclitaxel or Docetaxel) and Anthracycline regimens (followed by endocrine therapy for ER positive patients) and underwent surgery. Unfortunately, only a subset of this cohort had information on which Taxane each patient received, and therefore we chose to analyse both Taxane regimens together. All but 35 patients in the Hatzis cohort received their Taxane therapy prior to surgery, with exact timing of Taxane therapy per-patient was not recorded in the available clinical data. However, a 310 patient “discovery” sub-cohort was reported to have received all chemotherapy neoadjuvantly. Both these datasets had a pCR rate of 20%. Only *ODC1* and *EIF5A* were consistently measured on the microarrays used in the clinical studies. While *EIF5A* showed no meaningful correlation with pCR (Supplementary Figure 16a), *ODC1* was found to be predictive in both cohorts, with a AUROC of 0.73 in the MAQCII dataset, and AUROC of 0.71 in the Hatzis dataset respectively (AUPR of 0.44 and 0.33) (Figure 4a, 4b, Supplementary Figure 16b,c). Subsetting the Hatzis dataset to the “discovery” cohort, we found similar results (AUROC: 0.7, AUPR: 0.31, Supplementary Figure 16d,e). As such, all subsequent analysis was done on the full cohort. As expected, *ODC1* was found to be significantly higher expressed in responders compared to non-responders in both cohorts (Supplementary Figure 17).

**Fig. 4:**
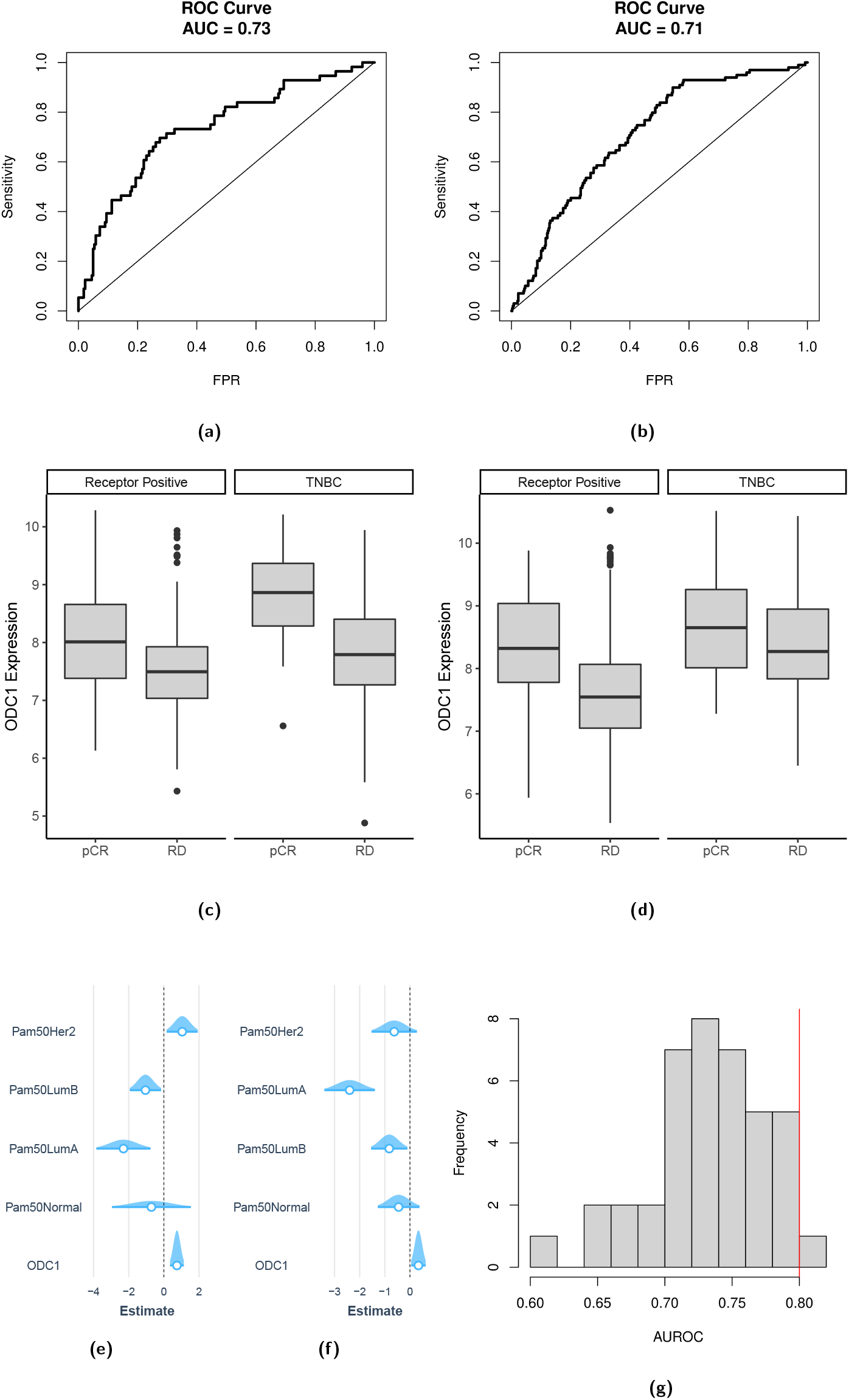
**a), b)** ROC curve for ODC1 expression as a predictor of pCR in the MAQCII and Hatzis datasets respectively; **c), d)** Expression of ODC1 stratified by TNBC status in the MAQCII and Hatzis datasets respectively; **e), f)** Coefficients and confidence intervals for logistic models predicting pCR from ODC1 expression and PAM50 subtypes for the MAQCII and Hatzis datasets respectively; **g)** the distribution of AUROC for predictive models trained by the MAQCII study authors as evaluated on the validation sub-cohort. The AUROC for ODC1 is marked by the red line.

*ODC1* has been reported in the literature to be overexpressed in the context of Triple Negative Breast Cancer [78]. Therefore, we examined whether the predictive power of *ODC1* could be explained by different response rates between TNBC and receptor positive cancers. We first verified that *ODC1* was correlated with TNBC status in these cohorts (Supplementary Figure 18). We then stratified patients by receptor status into TNBC and receptor positive sub-cohorts for both studies. In both studies, expression of *ODC1* was significantly higher in both subcohorts among responders compared to non-responders, suggesting that while *ODC1* is correlated with TNBC status, high *ODC1* expression can select for favourable response even after patient subtyping (Figure 4c, 4d). As IHC-determined subtypes within breast cancer are known to differ from expression-derived subtypes, we further also evaluated whether subtyping by gene expression, rather than receptor IHC status, would explain *ODC1* ‘s correlation with pCR. Using PAM50 classifications as published by the original study authors for the Hatzis dataset, we fit a logistic regression model to predict pCR from *ODC1* expression while adjusting for subtype. We found that even after adjustment for patient subtype, *ODC1* was found to be significantly predictive of response (*χ*^2^ test of likelihood p value: 0.021, Figure 4f).

For the MAQCII dataset, subtype labels were unavailable in the original study. Therefore, we used the `genefu` R package to predict PAM50 classification. We first verified the agreement between PAM50 labels for the Hatzis dataset generated by `genefu` with those published by the study authors, and found good concordance for all but the HER2+ subtype, which is unsurprising as the dataset selected for HER2 negative patients (Supplementary Table XXX). We then predicted PAM50 labels for each patient in the MAQCII dataset, and used them to fit a similar logistic regression model. Once again, *ODC1* was found to be predictive of response after adjusting for subtype (*χ*^2^ test of likelihood p value: 7.68·10^−5^, Figure 4e). Taken together, these results suggest that while BRCA subtypes affect the baseline expression of *ODC1, ODC1* is an independent predictor of response for neoadjuvant paclitaxel treatment.

The data for both BRCA cohorts examined in our study were originally collected to train and validate multivariate predictors for pCR response directly on clinical data. A natural question is to ask how *ODC1* performs as a predictor compared to multivariate models trained directly on these datasets. The authors of the MAQCII study trained and validated 40 different modeling approaches using an 80-20% split of their dataset for three different tasks, two of which were predicting pCR in an unselected, and ER-selected cohort of their study. Fortunately, they reported validation performance as AUROC for all their models, allowing us to compare their predictive power to the expression of *ODC1* as a biomarkers. On the 20% validation dataset, *ODC1* achieved an AUROC of 0.8 (Supplementary Figure 19a) for predicting pCR across all patients. Interestingly, only 1 of the 40 reported models outperformed *ODC1* as a predictor for this task (Figure 4g). Furthermore, when restricted to the harder task of predicting among ER-patients, *ODC1* outperformed all reported models in the study, with an AUROC of 0.71 vs a best AUROC of 0.627 for the multivariate models (Supplementary Figure 19b). It is important to note that predictors for pCR among ER-patients were trained only on ER-patients in the training set, greatly reducing the sample size available to the authors. This highlights the ability of robust, single gene markers to function as predictors in cases where more complex models fail due to a lack of training data.

## Discussion

Our study is the first systematic statistical meta-analysis of gene expression biomarkers of drug response across published *in vitro* pharmacogenomic datasets. We build upon previous work in statistics and econometrics literature to develop a three step, resampling-based inference approach to overcome violations of the assumptions behind standard asymptotic inference for meta-analysis. As our results revealed, approximately half the associations discovered in each individual study are not significant after meta-analysis. This coincides with previous work suggesting that differences in experimental protocols can drive large disagreements in drug response, which ultimately affect estimates of association with molecular features [27, 30, 32, 79]. In particular, differences in media conditions, experiment duration, cell density, and assay used to assess cell viability have all been previously shown to affect drug response measurements both due to changes attributable to technical factors (such as a failure to control for differences in cell growth rate), as well as by inducing biological variation among the same cell lines [30, 79]. This variation often affects measurements of response to different compounds to a different extent [30]. By applying a meta-analysis approach, our method implicitly favours those drugs and biomarkers which are less sensitive to these sources of variability across studies. While missing some associations between gene expression and drug response may be considered a limitation in the context of *in vitro* mechanistic studies, we consider robustness to experimental conditions crucial for translation of findings to the clinical context. Indeed, it would be surprising if a biomarker which failed to reproduce across different *in vitro* conditions would nevertheless be found to be consistently associated with response in the less controlled clinical context.

While our study is the first to apply a meta-analysis framework directly to *in vitro* biomarker discovery, existing work has used statistical methods to integrate data across studies. Previously, Jaiswal et al. [80] conducted a meta-analysis of preclinical pharmacogenomics studies with the goal of identifying novel cancer driver genes and therapeutic vulnerabilities. The rank-based method introduced by Jaiswal et al., termed CLIP, was focused on finding cell-line specific outlier molecular features that were supported by multiple datasets and multiple data modalities. To this end, drug response data was first transformed into gene-specific Target Addiction Scores [81]. As such, these data were used primarily to estimate the vulnerability of a cancer cell line to inhibition of particular targets, and not as a phenotype with which other molecular features were correlated. Our focus, on the other hand, is on discovering biomarkers for drug response directly. Furthermore, we focus on immediate translation, where the complexity of measuring multiple modalities is a liability to implementing findings into clinical practice. univariable transcriptomic biomarkers, although limited, have the potential for immediate translation into clinical practice, where univariable mutation-based features are already routinely accepted in the context of precision oncology research [1, 4, 5, 26].

In our study, we were able to confirm that the common practice of reducing the explored feature space to sets of genes previously known to be related to cancer, or otherwise well characterized, does in fact enrich for genes significantly associated with drug response. However, we observed that these cancer-related genesets are not necessarily a guarantee of consistency across studies. Often, reduced feature spaces are used for analysis when due to small sample sizes, multiple testing correction for the full transcriptome would reduce the power of the study to detect even the strongest expected effects, or when it is necessary to limit the capacity of multivariate models fit to the data. Our results largely support using cancer-related genesets to do such a priori feature selection, but highlight that the increase in statistical power will not necessarily lead to better reproducibility across studies. This aligns with previous research suggesting that differences in experimental conditions largely drives disagreements between studies, and not a lack of statistical power.

We also examined whether biomarkers which can be functionally linked to annotated drug targets tended to be better predictors of drug response than those without a known association with the drug mechanism of action. We found that neither distance in a Reactome pathway-generated gene network, nor whether the biomarker significantly correlated with CRISPR or RNAi-based inhibition of annotated drug targets predicted a stronger correlation with drug response, apart from the case where the expression of the target itself was a biomarker. A similar question was investigated by Gonçalves et al. [53], where “robust biomarkers” -biomarkers which predict both drug response and essentiality of genes which correlate with response to this drug - were hypothesized to be particularly promising candidates for translation. In their discussion, they suggest that by focusing on drugs with a significant association with gene knockdown and a shared biomarker for the knockdown and drug response may both select for drugs with greater specificity and increase the confidence in the discovered biomarkers - both desirable characteristics during preclinical drug development. Our study takes a different perspective, focusing on direct translation of existing preclinical results. Unfortunately, many currently used cancer therapies have complex mechanisms of action and are broad acting, which may preclude them from having strong associations with inhibition of any single gene. Supporting this hypothesis, Gonçalves et al. observed that drugs which had no strong correlation with CRISPR gene knockout profiles had, on average, significantly more drug targets [53]. As a concrete example, no CRISPR gene-knockout was found to significantly correlated with Paclitaxel in their study, which would preclude discovering a “robust biomarker” for this compound [53]. Furthermore, our study found no evidence that distance from the drug target in a Reactome pathway-derived network, nor correlation of a biomarker with CRISPR/RNAi inhibition of a drug target, would predict which biomarkers are reproducible across studies. We therefore focused our work purely on the robustness of biomarkers across independent studies, without requiring a notion of robustness across different technologies to inhibit particular drug targets.

Our pipeline detected the most associations among lung cancer cell lines out of all tissue types, with 87% of the significant associations discovered among this cancer type. We were able to identify two clusters of drugs with a large number of biomarkers predicting response to one cluster together with resistance to the second cluster. This clustering could at least partially be explained by differences in response between cells of a neuroendocrine vs alveolar epithelial origin. Neuroendocrine lung cancers are known to make up the majority of Small Cell Lung Cancers [59], while Non-Small Cell Lung Cancers are usually of other epithelial cell types. Interestingly, compounds with the same mechanisms as drugs from the first cluster are currently being investigated for their efficacy as novel therapeutics in SCLC. Navitoclax has been evaluated in a Phase I clinical trial for SCLC with promising results [61], while HDAC inhibitors have shown promising results in *in vivo* models of SCLC tumours[62, 82]. For both drug classes, markers of neuroendocrine cell state have been associated with better response to treatment [83, 84]. While histologically-identified SCLC is traditionally identified as a neuroendocrine tumour, there is an increasing acceptance of further subtyping of SCLC using molecular markers [59]. Upon subtyping, approximately 19% of SCLCs are found to be of subtypes which lack expression of neurendocrine cell markers [59]. Our results further suggest that transcriptomic expression of biomarkers is a better predictor of response than histopathological labeling of SCLC on its own, supporting the argument that these compounds should be investigated for their efficacy in a biomarker-selected population of patients.

Our meta-analysis is built on top of the work by the authors of all the pharmacogenomic studies that have published their data for researchers to use in their own research. These data, together with the work done previously by our group in reprocessing and standardizing public pharmacogenomics datasets [44, 85, 86] allowed our *in vitro* analysis to cover over 400 different drug-tissue combinations and evaluate millions of possible biomarkers. As our results demonstrate, while *in vitro* evidence can create hypotheses for predictive markers, many will not translate into clinically predictive features, with most of the biomarkers which we could evaluate in clinical data failing to predict drug response significantly better than random. Unfortunately, evaluating the translation potential of our findings was limited by the available clinical data for drug response with matched transcriptomic profiling of patients. The effort of CTRDB [33] to create a resource of well-annotated and uniformly processed clinical transcriptomic datasets is a crucial step in building the data resources needed to enable efficient translational research in predictive biomarker discovery. However, such resources can only work with the data which are collected and published. Many of the drug-tissue pairings we can explore *in vitro* are not being routinely used or currently investigated in the clinic and therefore clinical data cannot exist. However, 13 of the drug-tissue pairings for which we found significant biomarkers were cases where the drug was indicated for treatment of patients with cancers originating from that tissue type. Yet, only 5 of these 13 had data combining transcriptomics and clinical therapy response available. As evidence builds for the importance of expression information for precision oncology, we hope that transcriptomic profiling of patients becomes a more routine part of both clinical trial design as well as molecular characterization of biobanked samples or registry projects such as the AACR GENIE [5].

Rich clinical data combined with transcriptomic profiling was available for breast cancer patients treated with paclitaxel in combination with other chemotherapies in a neoadjuvant setting. Our preclinical meta-analysis found three significant markers of response to paclitaxel among breast cell lines: expression of the genes *ODC1, EIF4A1* and *EIF5A* was significantly associated sensitivity. We found that *ODC1* expression was predictive of response in two large breast cancer cohorts. This association with response remained after adjusting for IHC or expression derived subtyping of the tumours. ODC1 is the rate limiting enzyme of the polyamine synthesis pathway, with the main products being spermidine and spermine. Interestingly, EIF5A codes for the eukaryotic translation initiation factor 5A protein, which is the only human protein known to contain the uncommon amino acid hypusine [87]. Hypusine is essential to the function of EIF5A, and hypusine is synthesized in human cells from spermidine [87]. While the concurrent association of *ODC1* and *EIF5A* expression with response to paclitaxel is suggestive of a mechanistic relationship, experimental perturbation of the action of each protein in isolation would be required to determine whether this relationship is causal or simply correlative in nature.

The literature contains mixed evidence relating ODC1 expression or activity to response to chemotherapy. A previous *in vitro* study by Geck et al. in BRCA cell lines suggested that ODC1 inhibition can increase sensitivity to other chemotherapeutics [78], while work in HL-60 cells showed evidence that overexpression of ODC1 led to reduced Paclitaxel-induced apoptosis, and reduced Paclitaxel induced stalling in G2/M transition [88]. However, earlier work by Das et al. in the MCF7 cell line reported the inhibition of ODC1 activity to cause resistance to paclitaxel-induced cell death [89]. Their observations suggest a mechanism involving inhibition of paclitaxel induced apoptosis [89]. This coincides with an increasing understanding that different cellular contexts and different insults leading to apoptosis may affect whether ODC1 activity is pro- or anti-apoptotic [90]. Coinciding with this mechanistic evidence, Miolo et al. have previously identified circulating spermidine as a metabolic marker of response to Traztuzumab-Paclitaxel treatement in the context of neoadjuvant therapy of HER2 positive BRCA patients [91]. Taken in this context, our results suggest that clinical investigation of *ODC1* expression as a biomarker of neoadjuvant response to Paclitaxel among BRCA patients is warranted.

Overall, our work demonstrates how meta-analysis across pharmacogenomic studies can identify reproducible gene expression biomarkers. Whether a biomarker could be functionally linked to the mechanism of action of the drug was not a good predictor of neither the biomarker strength, nor of whether it would reproduce across datasets. We showed how transcriptomic biomarkers can predict differences in drug response among lung cancers better than disease type labels derived from histology. We were able to demonstrate an example of a preclinical biomarker which translates to a predictive marker in clinical data - expression of *ODC1* was predictive of paclitaxel response among BRCA patients. However, the majority of our results could not be clinically evaluated, as retrospective data for the drug-tissue combinations does not exist. We therefore intend our study both as a demonstration of the importance of proper meta-analysis of pharmacogenomic studies when doing biomarker discovery, as well as a resource of putative biomarkers with the best available *in vitro* supporting data for generating hypotheses and informing future translational research. This resource will be especially valuable as precision medicine trials adopt whole genome and transcriptome approaches to match patients to therapy, and these trials will in turn generate the data necessary to evaluate more of our discovered markers in a clinical context. Ultimately, it is our hope that comprehensive molecular profiling paired with meta-analysis of preclinical evidence becomes a routine source of information available to oncologists making treatment decisions.

## Methods

### Cell Line Dataset Access

All datasets were downloaded from the Orcestra platform [42] as PharmacoSet objects using the PharmacoGx R package [85].

### Transcriptomic Data Processsing

### Cell Line RNA Sequencing Data Processing

RNA sequencing data from cancer cell lines originating was processed as described previously [92]. In short, FASTQ files for the CCLE [41], gCSI [11, 17], GRAY [18], and UHNBreast [15] datasets were downloaded from original study authors. Kallisto (version 0.46.1) [93] was used to quantify gene expression as TPM values, using the GENCODE v33 transcriptome as reference. For meta-analysis, genes were filtered to protein coding genes as annotated by GENCODE v33.

### Cell Line Microarray Data Processing

CEL files for Affymetrix transcriptomic microarrays were downloaded from the repositories set up by the authors of the GDSC [14] and CCLE [9] studies. GDSC profiled cell lines on two independent microarrays platforms, the HT-HGU133A and the HGU219, and data from both were used in the study, as described in Collection of a compendium of preclinical cell line screening datasets. Microarray data were RMA [94] normalized within each dataset use the `affy` package [95] with the BrainArray CDF references (v 20.0.0) [96].

### Clinical Microarray Data Processing

Similar to the cell line microarray data, Affymetrix CEL files were downloaded from GEO, and RMA normalized using the `affy` package and the corresponding BrainArray CDF files (v 24.0.0).

### Processing of drug response data

Drug response data was downloaded from study-specific repositories and processed to quantify drug sensitivity as Area Above the Curve (AAC) values as previously described [32, 43, 44]. In summary, for the GDSC, gCSI, GRAY, and UHNBreast studies, well-level measurements were converted into % viability measurements by taking the average (median, unless more than 5 technical replicates were measured, then mean) of technical replicates for both treatment and control wells, subtracting the average plate specific background measurements, and taking the ratio between the treatment signal and control as a percentage. For the CTRPv2 and CCLE datasets, viability values were analyzed as published by the study authors. For each biological replicate of a drug-cell pairing in each dataset, 3-parameter Hill Curve models were fit to the dose-viability measurements using the PharmacoGx package [85]. Area Above the Curve values were calculated by taking the integral of the area over the measured concentration ranges, normalized by the maximum possible area over that concentration range. Prior to analyses, AAC values were averaged across biological replicates for each cell line - drug pair, if any existed within each dataset.

### QC and Filtering of Drug Dose Response Measurements

To ensure that AACs are comparable between each other within a single drug and dataset, we first standardized the measured concentration range across cell lines. For each drug within each dataset, we optimized a concentration range to retain the maximum number of measurements, while removing all experiments that did not cover the full range, and removing any points from experiments which lay outside this range. For all datasets, this removed less than 5% of the measured values. We then refit the curves and recalculated AAC values for each experiment. Finally, we filtered out dose-response curves which showed evidence of technical artifacts using our previously described `filterNoisyCurves` method [32]. This method filters out curves which show unexpectedly large oscillations between neighbouring points, or are significantly increasing over subsequent concentrations, giving evidence of drug dosing or labeling errors.

### Meta-Analysis Pipeline

Meta-analysis was conducted on the level of drug-gene-tissue triplets. As described above, genes were filtered to protein coding genes prior to analysis. Within each dataset, we identified those drug-gene-tissue triplets which had both drug response and gene expression measurements in at least 20 cell lines from that tissue type. We then kept all those triplets which had sufficient data in at least 3 studies. Data from studies with less than 20 samples for a drug-gene-tissue triplet was not considered.

No batch effect correction was done across datasets. Rather, the pipeline combines evidence on the level of correlation coefficients to deal with systemic offsets or differences in scale between studies.

### Within-Dataset Genewise Testing for Significance

Within each dataset, each drug-gene-tissue triplet was assessed for the presence of a significant correlation between the expression of the gene (either as TPM normalized RNAseq or RMA normalized microarray measurements) and the response to that drug (as AAC values) across cell lines from the tissue type. The correlation was measured using the Pearson Correlation Coefficient, and signficance testing was done using permutation testing as implemented by the `PharmacoGx` package [85], using the QUICK-STOP algorithm for early stopping [47], as described previously [44, 45]. Significance was assessed at an *α* = 0.05, after Bonferroni correction for multiple testing of all the genes being evaluated as possible biomarkers for the particular drug-tissue combination within each dataset. 95% Confidence intervals were evaluated using BCa-corrected bootstrap confidence intervals, with 1000 bootstrap samples per drug-gene-tissue triplet.

### Testing for Heterogeneity of Effects Across Studies

For each drug-gene-tissue triplet with at least 1 significant correlation detected across any of the datasets where if could be evaluated, we proceeded to perform meta-analysis across datasets. As the first step in meta-analysis, we evaluated whether there was evidence of heterogeneity across datasets. We first standardized the gene expression and AAC values within each dataset, and fit the following random-effect model to the data using `lme4` and REML estimation [97]:

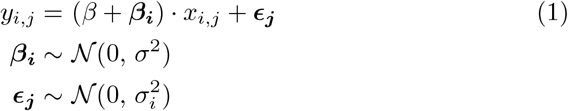

where *i* indexes the different studies, and *j* indexes samples within that study, *x* corresponds to standardized expression measurements, and *y* corresponds to standardized AAC measurements.

We then implemented the permutation test for the presence of a random effect across studies described by Lee and Braun [48]. In short, the method generates a permutation distribution *σ*^*\**^ for the random effect variance by permuting the errors *E* estimated by the residuals of (1), and then refitting the model to the simulated data. The significance is then assessed by calculating the proportion of (*σ*^*\**^)^2^ *> σ*^2^. For our study, we used an *α* = 0.1 to detect a significant presence of a random effect, as in this case Type II errors would lead to the choice of a more restrictive model, increasing the risk of Type I errors when assessing significance of the meta-estimates in the next step.

### Fixed and Random Effect Modeling Across Studies

For drug-gene-tissue triplets which did not show significant heterogeneity across studies, a fixed-effect model was used to compute a meta-estimate across studies. The meta-estimate for the Pearson Correlation Coefficient was computed by first standardizing the measured gene-expression and drug response values within each dataset, and then computing the regression coefficient between the standardized values across all datasets together. Significance was assessed using a permutation test, where prior to standardization within each dataset, the drug response was permuted across all measured values. The permuted values were standardized within each dataset, and a permuted Pearson correlation was computed. Two sided significance was evaluated by calculating the proportion of permuted correlations which had a larger deviation from zero compared to the observed meta-estimate. An *α* = 0.05 was used after Bonferroni correction for all the genes considered in our study across all drug-tissue combinations. The permutation testing was implemented as a C function called through the R interpreter.

For drug-gene-tissue triplets which did show significant heterogeneity across studies, a two-level, random-effect model was used to compute the meta-estimate for the correlation across studies. Similar to the fixed effect model, the gene expression and drug response values were first standardized within each study. The `MixedModels.jl` package in Julia was then used to estimate the two-level linear model described in (1) to the standardized data using REML estimation. The *β* parameter was then interpreted as the meta-estimate of the Pearson Correlation across studies, as it was the average/fixed regression coefficient estimated by the model when fit to standardized data.

Significance testing for the meta-estimate was done by testing the difference of *β* from 0. This was not possible to do with a permutation test, as under the permutation null, all studies have *β*_*i*_ = 0 in expectation. Therefore, a bootstrapping method was used to estimate significance. Bootstrap samples were generated through two-level sampling from unstandardized data: first, studies were resampled with replacement, then, for each resampled dataset, observations were resampled with replacement, in both cases sampling as many datasets/observations as there was in the original dataset. The data was then standardized within each dataset, and (1) was refit to each bootstrap sample. 1e7 bootstrap samples of *β*^*\**^ were calculated for each tissue-gene-drug triplet under analysis. The duality of confidence intervals and p-values [98] was used to calculate significance, by optimizing the largest *α* for which the percentile confidence interval estimated from the bootstrap distribution of *β*^*\**^ did not cross 0, and taking *p* = 1 − *α*.

95% Confidence intervals on the fixed effect estimate of the correlation from both fixed effect and random effect models were also evaluated using percentile bootstrap confidence intervals and the two-level resampling procedure described above, with 10,000 bootstrap samples used for the fixed effect model, and all 1e7 bootstrap samples used for the random-effect models.

Bias corrected and accelerated confidence intervals were not used for the meta-analysis models due to a limitation on calculating the acceleration correction. The resampling procedure did not draw samples according to an exponential-family of distributions (the total sample size varied across bootstrap samples, according to which datasets were resampled in the first stage). The efficient estimation of the acceleration correction presented by Efron from the empirical influence function assumes the data is resampled according to an exponential family [99], which is violated by our Data Generating Procedure. Note that previous simulation studies support the conjecture that BCa intervals would fail to show asymptotic refinement for two-level linear models [100].

### Biomarker Enrichment Analysis Among Cancer-Related Genesets

Enrichment analysis for biomarkers among Cancer Gene Census Genes, Cancer Functional Events, and Hallmark Pathways were done using the `fisher.test` function as implemented in R version 4.2.1. For testing over-representation of significant biomarkers among these genesets, the universe of genes considered was the intersection of all genes considered in the meta-analysis with all protein coding genes that were unique mappable to gene symbols using the Gencode v33 genome annotation. For testing overrepresentation of markers validating in meta-analysis among those which came up significant in single datasets, the universe considered was the union of all gene which were significant in at least 1 dataset.

The MSigDB Hallmark genesets v7.5.1 were used for this analysis. The Cancer Gene Census was accessed from https://cancer.sanger.ac.uk/census on March 22, 2021.

### Identification of Annotated Drug Targets

Drug target information was retrieved from Drugbank (v5.1.4), ChEMBL (file name: chembl drugtargets-19 _1_23_20), the CTRPv2 dataset (v20) and clue.io (dated 2018-09-07), and mapped to the drugs used within this study by matching (case insensitive) upon the compound names.

### Construction of Reactome Pathway Network and Testing of Association Between Distance to Target and Biomarker Strength

The gene pathway co-membership network was constructed according to the lowest level of the pathway hierarchy according to Reactome v79 [101], as accessed January 28, 2021. A gene-gene undirected graph was constructed by assigning based on co-membership in a human Reactome pathway. To calculate the distribution of distances between biomarkers and drug targets, for each biomarker (drug-gene-tissue triplet) significant after meta-analysis, the distance between the biomarker and the closest drug target in the network was calculated. For the distribution between drug targets and any random gene, for each drug, we randomly sampled the same number of genes from the Reactome network as there were biomarker for this drug, and calculated the minimum distance between each of these random genes and a drug target. For distances between pairs of random genes, for each drug we randomly sampled genes to create random drug targets and biomarker lists of the same length as the real targets/biomarkers. We then calculated the distance from each randomly sampled marker to the closest randomly sampled target. Significant differences between the distributions were tested using Mann-Whitney’s U test.

### Testing for Association of Discovered Diomarkers with Target Inhibition using DepMap CRISPR and RNAi Screens

Data for CRISPR gene deletion depletion effects as well as RNAi gene expression inhibition effects were downloaded from the DepMap data portal (version: 20Q2). For CRISPR, the `CRISPR_gene_effect.csv` file combining Achilles and Project SCORE data to estimate the effect on fitness from CRISPR deletion using the CERES pipeline [102] was used. For RNAi the `D2_combined_gene_dep_scores.csv` file combining the Marcotte et al. [15], Project DRIVE [57] and Achilles [54] datasets estimating the effect on cell line fitness of RNAi of each gene using the DEMETER2 pipeline [103] was used.

To discover biomarkers for CRISPR and RNAi scores, we considered the union of RNAseq gene expression data from the CCLE and GDSC, both processed as the PSet RNAseq datasets described above. When cell lines were profiled both in CCLE and GDSC, average expression for each gene was used. For each drug-gene-tissue biomarker triplet significant in at least one dataset (before meta-analysis), we tested the significance of the Pearson correlation between expression of the biomarker and CRISPR/RNAi scores of all annotated drug targets using the same early-stopping permutation method as described above in Within-Dataset Genewise Testing for Significance. Multiple testing correction was only applied for the total number of dataset-level biomarkers for each drug.

When examining the association between whether a biomarker predicts CRISPR/RNAi and distance to target in the Reactome network, or biomarker strength, only biomarkers significant after meta-analysis were considered.

### Clustering of Lung Cancer Biomarkers and Geneset Enrichment Analysis

#### Clustering of Drug by Lung Cancer Biomarker Associations

Interested in characterizing those biological processes that were consistently associated with sensitivity or resistance with several drugs, for this analyses we selected those drugs which had ¿=100 significant biomarkers, and those biomarkers which were associated with at least 3 drugs. This selected 324 genes for this analysis. To cluster the drugs by biomarker associations, we used the continuous correlation value for all significant associations, and a value of 0 for any gene that was not a biomarker for a particular drug. We then used a correlation distance between these biomarker vectors to perform Hierarchical clustering with Ward D2 linkage as implemented by R. Visual inspection of the clustering revealed a strong grouping into two equal clusters, which were termed Cluster 1 and Cluster 2 in the Results. We then assigned 316 of the 324 genes considered in this analysis to genesets predicting Cluster 1 or Cluster 2 sensitivities by assigning those genes for which the direction of correlation with drug response was consistent across all 4 drugs in each cluster.

#### GO Term Enrichment Analysis and EnrichmentMap Analysis

GO Biological Processes genesets were downloaded from MSigDB (C5 gene-sets, version 7.1). Overrepesentation analysis was done using the `fora` function from the `fgsea` package [104]. The universe considered were all protein coding genes mappable to a gene symbol in the GENCODE annotation (v33). Overrepresentation analysis was done separately for the genes associated with Cluster 1 and Cluster 2. The results of this analyses were then assigned opposite enrichment directions and read into the EnrichmentMap Cytoscape app [105]. A GO term network was created by examining gene co-membership between GO terms that were enriched among the Cluster 1 or Cluster 2 genes (with q value ¡= 0.05), with a minimum overlap between genesets of 0.375 Jaccard. These GO Terms were then clustered using the Markov Chain Clustering algorithm as implemented by the `clusterMaker` Cytoscape app, with default parameters other than setting the overlap to be used as the edge weight during clustering. The top ten clusters by GO term membership were then analyzed for commonly (co-)occurring terms within their descriptions using the `AutoAnnotate` app to derive 3-word labels for each cluster derived form the commonly observed descriptors.

#### Enrichment of Cell Type Markers

Cell Type Marker Genesets were downloaded from MSigDB (C8 genesets, version 7.5.1) and subset to only those genesets identified from single cell profiling by Travaglini et al [60]. Overrepesentation analysis was done as in GO Term Enrichment Analysis and EnrichmentMap Analysis.

### Binarization of YAP1 and INSM1 Expression and Comparison to Disease Labels

To assign high/low labels for expression of YAP1 and INSM1 to each lung cancer cell line within each dataset, a mixture of two Gaussians was fit to the distribution of gene expression with the datasets as described before [106]. High-low labels were then assigned by assigning each cell line to the higher or lower mode in the data according to maximum posterior probability fit by the model.

Cell lines were also classified to Small Cell Lung Cancer or Non-Small Cell Lung Cancer groups by mapping the disease annotation of each cell line in the Cellosaurus database [107] to the OncoTree Cancer Ontology [108] as described previously [44].

The Area Under the Receiver Operating Characteristic curve (AUROC) between the AAC as a continuous number for each drug of interest and the Disease or Expression labels were computed and the distribution across drugs was compared using a Wilcoxon Signed-Rank Test.

### Testing of Significance of Gene Expression Biomarker Associations with Response in Clinical Datasets

To assess whether *in vitro* discovered biomarkers are predictive of response clinically, we evaluated whether gene expression was significantly associated with response annotations among several clinical trial datasets. We aimed to assess response in the same way as the original study authors, relying on the published clinical data metadata. The data from the COG trials 8704 and 9404 were downloaded from GEO datasets GSE14613 and GSE14615. Response was evaluated in Complete Continuous Response, Relapsed, and Induction Failure categories. Data for the Ovarian datasets were download from GEO datasets: GSE14764, GSE15622, GSE63885 and GSE31245. Measurements of residual disease during surgery were used as response labels for GSE14764. Response/Non-Response labels from GSE15622 based on CA125 measurements were used as published by study authors. While pre- and post-treatment samples were available in GSE15622, only pre-treatment samples were considered. RECIST-like response assessment was binarized into Responders (Complete Response and Partial Response) and Non-Responders (Stable Disease and Progressive Disease) for analysis of GSE63885. No response information was available for GSE31245, and biomarker validation was done through comparison with censored Overall Survival times. Data from the BATTLE1 trial were accessed from GSE33072. As the Erlotinib cohort had no recorded response measurements, biomarker expression was compared with Progression Free Survival. Data from the CHER-LOB study were downloaded from GSE66305. Response was evaluated as pathological Complete Response (pCR) vs Residual Disease as published by the authors. Similarly, data for the MAQCII and Hatzis datasets were downloaded from GSE20194 and GSE25066, and in both cases response was evaluated as pCR vs RD labels from the original studies.

Statistical comparisons between gene expression and response for all binary, two-group measurements were done using the Wilcoxon rank-sum/Mann-Whitney U tests. 3 group analyses were done using the Kruskal-Walis test. Comparisons between expression and survival were done using a permutation tests on the Concordance Index. The predictive power of significant biomarkers was quantified as AUROC and AUPR measures.

## References

[1] Remon, J. & Dienstmann, R. Precision oncology: Separating the wheat from the chaff. ESMO Open 3 (6), e000446 (2018). https://doi.org/10.1136/esmoopen-2018-000446.

[2] Stockley, T. L. et al. Molecular profiling of advanced solid tumors and patient outcomes with genotype-matched clinical trials: The Princess Margaret IMPACT/COMPACT trial. Genome Med 8 (1), 109 (2016). https://doi.org/10.1186/s13073-016-0364-2.

[3] Tsimberidou, A.-M. et al. Initiative for Molecular Profiling and Advanced Cancer Therapy (IMPACT): An MD Anderson Precision Medicine Study. JCO Precision Oncology (1), 1–18 (2017). URL http://ascopubs.org/doi/abs/10.1200/PO.17.00002. https://doi.org/10.1200/PO.17.00002.

[4] Malone, E. R., Oliva, M., Sabatini, P. J. B., Stockley, T. L. & Siu, L. L. Molecular profiling for precision cancer therapies. Genome Medicine 12 (1), 8 (2020). URL https://doi.org/10.1186/s13073-019-0703-1. https://doi.org/10.1186/s13073-019-0703-1.

[5] Pugh, T. J. et al. AACR Project GENIE: 100,000 cases and beyond. Cancer Discovery cd.21.1547 (2022). URL https://doi.org/10.1158/2159-8290.CD-21-1547. https://doi.org/10.1158/2159-8290.CD-21-1547

[6] Tsimberidou, A. M., Fountzilas, E., Nikanjam, M. & Kurzrock, R. Review of precision cancer medicine: Evolution of the treatment paradigm. Cancer Treat Rev 86, 102019 (2020). https://doi.org/10.1016/j.ctrv.2020.102019.

[7] Chakravarty, D. et al. OncoKB: A Precision Oncology Knowledge Base. JCO Precision Oncology (1), 1–16 (2017). URL https://ascopubs.org/doi/full/10.1200/PO.17.00011. https://doi.org/10.1200/PO.17.00011.

[8] Leo, C. P., Leo, C. & Szucs, T. D. Breast cancer drug approvals by the US FDA from 1949 to 2018. Nature Reviews Drug Discovery 19 (1), 11–11 (2019). URL https://www.nature.com/articles/d41573-019-00201-w. https://doi.org/10.1038/d41573-019-00201-w.

[9] Barretina, J. et al. The Cancer Cell Line Encyclopedia enables predictive modelling of anticancer drug sensitivity. Nature 483 (7391), 603–607 (2012). URL http://www.nature.com.myaccess.library.utoronto.ca/nature/journal/v483/n7391/full/nature11003.html. https://doi.org/10.1038/nature11003.

[10] Seashore-Ludlow, B. et al. Harnessing Connectivity in a Large-Scale Small-Molecule Sensitivity Dataset. Cancer Discovery (2015). URL http://cancerdiscovery.aacrjournals.org/content/early/2015/10/14/2159-8290.CD-15-0235. https://doi.org/10.1158/2159-8290.CD-15-0235

[11] Haverty, P. M. et al. Reproducible pharmacogenomic profiling of cancer cell line panels. Nature 533 (7603), 333–337 (2016). URL https://www.nature.com/articles/nature17987. https://doi.org/10.1038/nature17987

[12] Iorio, F. et al. A Landscape of Pharmacogenomic Interactions in Cancer. Cell 166 (3), 740–754 (2016). URL http://www.cell.com/cell/abstract/S0092-8674(16)30746-2. https://doi.org/10.1016/j.cell.2016.06.017.

[13] Feizi, N. et al. PharmacoDB 2.0: Improving scalability and transparency of in vitro pharmacogenomics analysis. Nucleic Acids Research 50 (D1), D1348–D1357 (2022). URL https://doi.org/10.1093/nar/gkab1084. https://doi.org/10.1093/nar/gkab1084.

[14] Garnett, M. J. et al. Systematic identification of genomic markers of drug sensitivity in cancer cells. Nature 483 (7391), 570–575 (2012). URL http://www.nature.com/articles/nature11005. https://doi.org/10.1038/nature11005.

[15] Marcotte, R. et al. Functional Genomic Landscape of Human Breast Cancer Drivers, Vulnerabilities, and Resistance. Cell 164 (1), 293–309 (2016). URL http://www.cell.com/cell/abstract/S0092-8674(15)01624-4. https://doi.org/10.1016/j.cell.2015.11.062.

[16] Rees, M. G. et al. Correlating chemical sensitivity and basal gene expression reveals mechanism of action. Nat Chem Biol 12 (2), 109–116 (2016). URL http://www.nature.com/nchembio/journal/v12/n2/full/nchembio.1986.html. https://doi.org/10.1038/nchembio.1986.

[17] Klijn, C. et al. A comprehensive transcriptional portrait of human cancer cell lines. Nat Biotech 33 (3), 306–312 (2015). URL http://www.nature.com/nbt/journal/v33/n3/full/nbt.3080.html. https://doi.org/10.1038/nbt.3080.

[18] Daemen, A. et al. Modeling precision treatment of breast cancer. Genome Biology 14, R110 (2013). URL http://dx.doi.org/10.1186/gb-2013-14-10-r110. https://doi.org/10.1186/gb-2013-14-10-r110.

[19] Hafner, M. et al. Quantification of sensitivity and resistance of breast cancer cell lines to anti-cancer drugs using GR metrics. Sci Data 4 (2017). URL https://www.ncbi.nlm.nih.gov/pmc/articles/PMC5674849/. https://doi.org/10.1038/sdata.2017.166.

[20] Shoemaker, R. H. The NCI60 human tumour cell line anticancer drug screen. Nat Rev Cancer 6 (10), 813–823 (2006). URL https://www.nature.com/articles/nrc1951. https://doi.org/10.1038/nrc1951.

[21] Corsello, S. M. et al. Discovering the anticancer potential of non-oncology drugs by systematic viability profiling. Nat Cancer (2), 1–14 (2020). URL https://www.nature.com/articles/s43018-019-0018-6. https://doi.org/10.1038/s43018-019-0018-6.

[22] Costello, J. C. et al. A community effort to assess and improve drug sensitivity prediction algorithms. Nat Biotech 32 (12), 1202–1212 (2014). URL http://www.nature.com/nbt/journal/v32/n12/full/nbt.2877.html. https://doi.org/10.1038/nbt.2877.

[23] Roumeliotis, T. I. et al. Genomic Determinants of Protein Abundance Variation in Colorectal Cancer Cells. Cell Reports 20 (9), 2201–2214 (2017). URL http://www.cell.com/cell-reports/abstract/S2211-1247(17)31100-2. https://doi.org/10.1016/j.celrep.2017.08.010.

[24] Rodon, J. et al. Genomic and transcriptomic profiling expands precision cancer medicine: The WINTHER trial. Nat Med 25 (5), 751–758 (2019). URL https://www.nature.com/articles/s41591-019-0424-4. https://doi.org/10.1038/s41591-019-0424-4.

[25] Horak, P. et al. Comprehensive Genomic and Transcriptomic Analysis for Guiding Therapeutic Decisions in Patients with Rare Cancers. Cancer Discovery 11 (11), 2780–2795 (2021). URL https://doi.org/10.1158/2159-8290.CD-21-0126. https://doi.org/10.1158/2159-8290.CD-21-0126

[26] Pleasance, E. et al. Whole-genome and transcriptome analysis enhances precision cancer treatment options. Annals of Oncology (2022). URL https://www.sciencedirect.com/science/article/pii/S0923753422017239. https://doi.org/10.1016/j.annonc.2022.05.522.

[27] Haibe-Kains, B. et al. Inconsistency in large pharmacogenomic studies. Nature 504 (7480), 389–393 (2013). URL http://www.nature.com.myaccess.library.utoronto.ca/nature/journal/v504/n7480/full/nature12831.html. https://doi.org/10.1038/nature12831.

[28] Hafner, M., Niepel, M. & Sorger, P. K. Alternative drug sensitivity metrics improve preclinical cancer pharmacogenomics. Nature Biotechnology (2017). URL https://www.nature.com/articles/nbt.3882. https://doi.org/10.1038/nbt.3882.

[29] Safikhani, Z. et al. Revisiting inconsistency in large pharmacogenomic studies. F1000Research 5, 2333 (2017). URL https://f1000research.com/articles/5-2333/v3. https://doi.org/10.12688/f1000research.9611.3.

[30] Niepel, M. et al. A Multi-center Study on the Reproducibility of Drug-Response Assays in Mammalian Cell Lines. Cell Systems 9 (1), 35– 48.e5 (2019). URL https://www.sciencedirect.com/science/article/pii/S2405471219302005. https://doi.org/10.1016/j.cels.2019.06.005.

[31] Mpindi, J. P. et al. Consistency in drug response profiling. Nature 540 (7631), E5–E6 (2016). URL http://dx.doi.org/10.1038/nature20171

[32] Safikhani, Z. et al. Revisiting inconsistency in large pharmacogenomic studies. F1000Research 5 (2016).

[33] Liu, Z. et al. CTR-DB, an omnibus for patient-derived gene expression signatures correlated with cancer drug response. Nucleic Acids Research 50 (D1), D1184–D1199 (2022). URL https://doi.org/10.1093/nar/gkab860. https://doi.org/10.1093/nar/gkab860.

[34] Lee, J. S. et al. Synthetic lethality-mediated precision oncology via the tumor transcriptome. Cell 184 (9), 2487–2502.e13 (2021). URL https://www.sciencedirect.com/science/article/pii/S0092867421003615. https://doi.org/10.1016/j.cell.2021.03.030.

[35] Heiser, L. M. et al. Subtype and pathway specific responses to anticancer compounds in breast cancer. Proceedings of the National Academy of Sciences 109 (8), 2724–2729 (2012).

[36] Marcotte, R. et al. Essential Gene Profiles in Breast, Pancreatic, and Ovarian Cancer Cells. Cancer Discovery 2 (2), 172–189 (2012).

[37] Safikhani, Z. et al. Gene isoforms as expression-based biomarkers predictive of drug response in vitro. Nature Communications 8 (1), 1126 (2017). URL https://www.nature.com/articles/s41467-017-01153-8. https://doi.org/10.1038/s41467-017-01153-8.

[38] Yang, W. et al. Genomics of Drug Sensitivity in Cancer (GDSC): A resource for therapeutic biomarker discovery in cancer cells. Nucleic acids research 41 (D1), D955–D961 (2012).

[39] Picco, G. et al. Functional linkage of gene fusions to cancer cell fitness assessed by pharmacological and CRISPR-Cas9 screening. Nature Communications 10 (1), 2198 (2019). URL https://www.nature.com/articles/s41467-019-09940-1. https://doi.org/10.1038/s41467-019-09940-1.

[40] Basu, A. et al. An Interactive Resource to Identify Cancer Genetic and Lineage Dependencies Targeted by Small Molecules. Cell 154 (5), 1151–1161 (2013).

[41] Ghandi, M. et al. Next-generation characterization of the Cancer Cell Line Encyclopedia. Nature 569 (7757), 503–508 (2019). URL https://www.nature.com/articles/s41586-019-1186-3. https://doi.org/10.1038/s41586-019-1186-3.

[42] Mammoliti, A. et al. Orchestrating and sharing large multimodal data for transparent and reproducible research. Nat Commun 12 (1), 5797 (2021). URL https://www.nature.com/articles/s41467-021-25974-w. https://doi.org/10.1038/s41467-021-25974-w.

[43] Smirnov, P. et al. PharmacoDB: An integrative database for mining in vitro anticancer drug screening studies. Nucleic acids research 46 (D1), D994–D1002 (2018).

[44] Feizi, N. et al. PharmacoDB 2.0: Improving scalability and transparency of in vitro pharmacogenomics analysis. Nucleic acids research 50 (D1), D1348–D1357 (2022).

[45] Smirnov, P. et al. Evaluation of statistical approaches for association testing in noisy drug screening data. BMC Bioinformatics 23, 188 (2022). URL https://www.ncbi.nlm.nih.gov/pmc/articles/PMC9118710/. https://doi.org/10.1186/s12859-022-04693-z.

[46] Geeleher, P., Gamazon, E. R., Seoighe, C., Cox, N. J. & Huang, R. S. Consistency in large pharmacogenomic studies. Nature 540 (7631), E1–E2 (2016). URL http://www.nature.com/articles/nature19838. https://doi.org/10.1038/nature19838.

[47] Hecker, J. et al. A flexible and nearly optimal sequential testing approach to randomized testing: QUICK-STOP. Genet Epidemiol 44 (2), 139–147 (2020). https://doi.org/10.1002/gepi.22268.

[48] Lee, O. E. & Braun, T. M. Permutation Tests for Random Effects in Linear Mixed Models. Biometrics 68 (2) (2012). URL https://www.ncbi.nlm.nih.gov/pmc/articles/PMC3883440/. https://doi.org/10.1111/j.1541-0420.2011.01675.x.

[49] Sondka, Z. et al. The COSMIC Cancer Gene Census: Describing genetic dysfunction across all human cancers. Nat Rev Cancer 18 (11), 696–705 (2018). URL http://www.nature.com/articles/s41568-018-0060-1. https://doi.org/10.1038/s41568-018-0060-1.

[50] Subramanian, A. et al. Gene set enrichment analysis: A knowledge-based approach for interpreting genome-wide expression profiles. PNAS 102 (43), 15545–15550 (2005). URL http://www.pnas.org/content/102/43/15545. https://doi.org/10.1073/pnas.0506580102.

[51] Liberzon, A. et al. Molecular signatures database (MSigDB) 3.0. Bioinformatics 27 (12), 1739 (2011). URL https://www.ncbi.nlm.nih.gov/pmc/articles/PMC3106198/. https://doi.org/10.1093/bioinformatics/btr260.

[52] Liberzon, A. et al. The Molecular Signatures Database (MSigDB) hallmark gene set collection. Cell Syst 1 (6), 417–425 (2015). URL https://www.ncbi.nlm.nih.gov/pmc/articles/PMC4707969/. https://doi.org/10.1016/j.cels.2015.12.004.

[53] Gonçcalves, E. et al. Drug mechanism-of-action discovery through the integration of pharmacological and CRISPR screens. Molecular Systems Biology 16 (7) (2020). URL https://onlinelibrary.wiley.com/doi/10.15252/msb.20199405. https://doi.org/10.15252/msb.20199405.

[54] Tsherniak, A. et al. Defining a Cancer Dependency Map. Cell 170 (3), 564–576.e16 (2017). URL https://www.cell.com/cell/abstract/S0092-8674(17)30651-7. https://doi.org/10.1016/j.cell.2017.06.010.

[55] Dempster, J. M. et al. Agreement between two large pan-cancer CRISPR-Cas9 gene dependency data sets. Nat Commun 10 (1), 5817 (2019). https://doi.org/10.1038/s41467-019-13805-y.

[56] Behan, F. M. et al. Prioritization of cancer therapeutic targets using CRISPR-Cas9 screens. Nature 568 (7753), 511–516 (2019). https://doi.org/10.1038/s41586-019-1103-9.

[57] McDonald, E. R. et al. Project DRIVE: A Compendium of Cancer Dependencies and Synthetic Lethal Relationships Uncovered by Large-Scale, Deep RNAi Screening. Cell 170 (3), 577–592.e10 (2017). https://doi.org/10.1016/j.cell.2017.07.005.

[58] Hart, T. et al. High-Resolution CRISPR Screens Reveal Fitness Genes and Genotype-Specific Cancer Liabilities. Cell 163 (6), 1515–1526 (2015). https://doi.org/10.1016/j.cell.2015.11.015.

[59] Rudin, C. M. et al. Molecular subtypes of small cell lung cancer: A synthesis of human and mouse model data. Nat Rev Cancer 19 (5), 289–297 (2019). URL https://www.nature.com/articles/s41568-019-0133-9. https://doi.org/10.1038/s41568-019-0133-9.

[60] Travaglini, K. J. et al. A molecular cell atlas of the human lung from single-cell RNA sequencing. Nature 587 (7835), 619–625 (2020). URL http://www.nature.com/articles/s41586-020-2922-4. https://doi.org/10.1038/s41586-020-2922-4.

[61] Gandhi, L. et al. Phase I Study of Navitoclax (ABT-263), a Novel Bcl-2 Family Inhibitor, in Patients With Small-Cell Lung Cancer and Other Solid Tumors. JCO 29 (7), 909–916 (2011). URL https://ascopubs.org/doi/10.1200/JCO.2010.31.6208. https://doi.org/10.1200/JCO.2010.31.6208.

[62] Jia, D. et al. Crebbp Loss Drives Small Cell Lung Cancer and Increases Sensitivity to HDAC Inhibition. Cancer Discovery 8 (11), 1422–1437 (2018). URL https://doi.org/10.1158/2159-8290.CD-18-0385. https://doi.org/10.1158/2159-8290.CD-18-0385.

[63] Qin, A. et al. Trial in progress: A multicenter phase Ib/II study of pelcitoclax (APG-1252) in combination with paclitaxel in patients with relapsed/refractory small-cell lung cancer (R/R SCLC). JCO 39 (15 suppl), TPS8589–TPS8589 (2021). URL https://ascopubs.org/doi/abs/10.1200/JCO.2021.39.15suppl.TPS8589. https://doi.org/10.1200/JCO.2021.39.15suppl.TPS8589.

[64] Rudin, C. M. et al. A pilot trial of G3139, a bcl-2 antisense oligonucleotide,and paclitaxel in patients with chemorefractory small-cell lung cancer. Annals of Oncology 13 (4), 539–545 (2002). URL https://www.sciencedirect.com/science/article/pii/S0923753419618867. https://doi.org/10.1093/annonc/mdf124.

[65] Warren, A. et al. Global computational alignment of tumor and cell line transcriptional profiles. Nature Communications 12 (1), 22 (2021). URL https://www.nature.com/articles/s41467-020-20294-x. https://doi.org/10.1038/s41467-020-20294-x.

[66] Mourragui, S. et al. PRECISE+ predicts drug response in patients by non-linear subspace-based transfer from cell lines and PDX models. bioRxiv 2020.06.29.177139 (2020). URL https://www.biorxiv.org/content/10.1101/2020.06.29.177139v2. https://doi.org/10.1101/2020.06.29.177139.

[67] Mourragui, S. M. C. et al. Predicting patient response with models trained on cell lines and patient-derived xenografts by nonlinear transfer learning. PNAS 118 (49) (2021). URL https://www.pnas.org/content/118/49/e2106682118. https://doi.org/10.1073/pnas.2106682118.

[68] Anderson, T. W. & Darling, D. A. Asymptotic Theory of Certain “Goodness of Fit” Criteria Based on Stochastic Processes. The Annals of Mathematical Statistics 23 (2), 193–212 (1952). URL https://projecteuclid.org/journals/annals-of-mathematical-statistics/volume-23/issue-2/Asymptotic-Theory-of-Certain-Goodness-of-Fit-Criteria-Based-on/10.1214/aoms/1177729437.full. https://doi.org/10.1214/aoms/1177729437.

[69] Winter, S. S. et al. Identification of genomic classifiers that distinguish induction failure in T-lineage acute lymphoblastic leukemia: A report from the Children’s Oncology Group. Blood 110 (5), 1429–1438 (2007). URL https://doi.org/10.1182/blood-2006-12-059790. https://doi.org/10.1182/blood-2006-12-059790.

[70] Lisowska, K. M. et al. Gene Expression Analysis in Ovarian Cancer – Faults and Hints from DNA Microarray Study. Frontiers in Oncology 4 (2014).

[71] Denkert, C. et al. A prognostic gene expression index in ovarian cancer—validation across different independent data sets. The Journal of Pathology 218 (2), 273–280 (2009). URL http://onlinelibrary.wiley.com/doi/abs/10.1002/path.2547. https://doi.org/10.1002/path.2547.

[72] Ahmed, A. A. et al. The Extracellular Matrix Protein TGFBI Induces Microtubule Stabilization and Sensitizes Ovarian Cancers to Paclitaxel. Cancer Cell 12 (6), 514–527 (2007). URL https://www.ncbi.nlm.nih.gov/pmc/articles/PMC2148463/. https://doi.org/10.1016/j.ccr.2007.11.014.

[73] Spentzos, D. et al. Gene expression signature with independent prognostic significance in epithelial ovarian cancer. Journal of clinical oncology: official journal of the American Society of Clinical Oncology 22 (23), 4700–4710 (2004).

[74] Byers, L. A. et al. An epithelial-mesenchymal transition gene signature predicts resistance to EGFR and PI3K inhibitors and identifies Axl as a therapeutic target for overcoming EGFR inhibitor resistance. Clin Cancer Res 19 (1), 279–290 (2013). https://doi.org/10.1158/1078-0432.CCR-12-1558.

[75] Guarneri, V. et al. Prospective Biomarker Analysis of the Randomized CHER-LOB Study Evaluating the Dual Anti-HER2 Treatment With Trastuzumab and Lapatinib Plus Chemotherapy as Neoadjuvant Therapy for HER2-Positive Breast Cancer. Oncologist 20 (9), 1001–1010 (2015). https://doi.org/10.1634/theoncologist.2015-0138.

[76] Hatzis, C. et al. A genomic predictor of response and survival following taxane-anthracycline chemotherapy for invasive breast cancer. JAMA 305 (18), 1873–1881 (2011). https://doi.org/10.1001/jama.2011.593.

[77] Popovici, V. et al. Effect of training-sample size and classification difficulty on the accuracy of genomic predictors. Breast Cancer Res 12 (1), R5 (2010). URL https://www.ncbi.nlm.nih.gov/pmc/articles/PMC2880423/. https://doi.org/10.1186/bcr2468.

[78] Geck, R. C. et al. Inhibition of the polyamine synthesis enzyme ornithine decarboxylase sensitizes triple-negative breast cancer cells to cytotoxic chemotherapy. Journal of Biological Chemistry 295 (19), 6263–6277 (2020). URL https://www.sciencedirect.com/science/article/pii/S0021925817484936. https://doi.org/10.1074/jbc.RA119.012376.

[79] Hafner, M., Niepel, M., Chung, M. & Sorger, P. K. Growth rate inhibition metrics correct for confounders in measuring sensitivity to cancer drugs. Nature Methods 13 (6), 521–527 (2016). URL https://www.nature.com/articles/nmeth.3853. https://doi.org/10.1038/nmeth.3853.

[80] Jaiswal, A. et al. Multi-modal meta-analysis of cancer cell line omics profiles identifies ECHDC1 as a novel breast tumor suppressor. Molecular Systems Biology 17 (3), e9526 (2021). URL https://www.embopress.org/doi/10.15252/msb.20209526. https://doi.org/10.15252/msb.20209526.

[81] Jaiswal, A., Yadav, B., Wennerberg, K. & Aittokallio, T. Integrated Analysis of Drug Sensitivity and Selectivity to Predict Synergistic Drug Combinations and Target Coaddictions in Cancer. Methods Mol Biol 1888, 205–217 (2019). https://doi.org/10.1007/978-1-4939-8891-412.

[82] Crisanti, M. C. et al. The HDAC inhibitor panobinostat (LBH589) inhibits mesothelioma and lung cancer cells in vitro and in vivo with particular efficacy for small cell lung cancer. Molecular Cancer Therapeutics 8 (8), 2221–2231 (2009). URL https://doi.org/10.1158/1535-7163.MCT-09-0138. https://doi.org/10.1158/1535-7163.MCT-09-0138.

[83] Jia, D. et al. Crebbp Loss Drives Small Cell Lung Cancer and Increases Sensitivity to HDAC Inhibition. Cancer Discovery 8 (11), 1422–1437 (2018). URL https://doi.org/10.1158/2159-8290.CD-18-0385. https://doi.org/10.1158/2159-8290.CD-18-0385.

[84] Lochmann, T. L. et al. Venetoclax Is Effective in Small-Cell Lung Cancers with High BCL-2 Expression. Clinical Cancer Research 24 (2), 360–369 (2018). URL https://doi.org/10.1158/1078-0432.CCR-17-1606. https://doi.org/10.1158/1078-0432.CCR-17-1606.

[85] Smirnov, P. et al. PharmacoGx: An R package for analysis of large pharmacogenomic datasets. Bioinformatics 32 (8), 1244–1246 (2016). https://doi.org/10.1093/bioinformatics/btv723.

[86] Smirnov, P. et al. PharmacoDB: An integrative database for mining in vitro anticancer drug screening studies. Nucleic Acids Res 46 (D1), D994–D1002 (2018). URL https://academic.oup.com/nar/article/46/D1/D994/4372597. https://doi.org/10.1093/nar/gkx911.

[87] Park, M. H. & Wolff, E. C. Hypusine, a polyamine-derived amino acid critical for eukaryotic translation. J Biol Chem 293 (48), 18710–18718 (2018). URL https://www.ncbi.nlm.nih.gov/pmc/articles/PMC6290153/. https://doi.org/10.1074/jbc.TM118.003341.

[88] Hsu, P.-C. et al. Ornithine decarboxylase attenuates leukemic chemotherapy drugs-induced cell apoptosis and arrest in human promyelocytic HL-60 cells. Leukemia Research 32 (10), 1530–1540 (2008). URL https://www.sciencedirect.com/science/article/pii/S0145212608000568. https://doi.org/10.1016/j.leukres.2008.01.017.

[89] Das, B., Rao, A. R. & Madhubala, R. Difluoromethylornithine antagonizes taxol cytotoxicity in MCF-7 human breast cancer cells. Oncol Res 9 (11-12), 565–572 (1997).

[90] Seiler, N. & Raul, F. Polyamines and apoptosis. J Cell Mol Med 9 (3), 623–642 (2005 Jul-Sep). https://doi.org/10.1111/j.1582-4934.2005.tb00493.x.

[91] Miolo, G. et al. Pharmacometabolomics study identifies circulating spermidine and tryptophan as potential biomarkers associated with the complete pathological response to trastuzumab-paclitaxel neoadjuvant therapy in HER-2 positive breast cancer. Oncotarget 7 (26), 39809–39822 (2016). URL https://www.ncbi.nlm.nih.gov/pmc/articles/PMC5129972/. https://doi.org/10.18632/oncotarget.9489.

[92] Sharifi-Noghabi, H. et al. Drug sensitivity prediction from cell line-based pharmacogenomics data: Guidelines for developing machine learning models. Briefings in Bioinformatics 22 (6), bbab294 (2021).

[93] Bray, N. L., Pimentel, H., Melsted, P. & Pachter, L. Near-optimal probabilistic RNA-seq quantification. Nat Biotechnol 34 (5), 525–527 (2016). URL https://www.nature.com/articles/nbt.3519. https://doi.org/10.1038/nbt.3519.

[94] Irizarry, R. A. et al. Exploration, normalization, and summaries of high density oligonucleotide array probe level data. Biostatistics (Oxford, England) 4 (2), 249–264 (2003).

[95] Gautier, L., Cope, L. M., Bolstad, B. M. & Irizarry, R. A. Affy - Analysis of Affymetrix GeneChip data at the probe level. Bioinformatics 20 (3), 307–315 (2004).

[96] Dai, M. et al. Evolving gene/transcript definitions significantly alter the interpretation of GeneChip data. Nucl. Acids Res. 33 (20), e175.#x2013; (2005).

[97] Bates, D., Mächler, M., Bolker, B. & Walker, S. Fitting Linear Mixed-Effects Models Using lme4. J. Stat. Soft. 67 (1) (2015). URL http://www.jstatsoft.org/v67/i01/. https://doi.org/10.18637/jss.v067.i01.

[98] Efron, B. & Tibshirani, R. J. An Introduction to the Bootstrap (Chapman and Hall, New York, 1993).

[99] Efron, B. Better Bootstrap Confidence Intervals. Journal of the American Statistical Association 82 (397), 171–185 (1987). URL https://www.jstor.org/stable/2289144. https://doi.org/10.2307/2289144.

[100] Cameron, A. C., Gelbach, J. B. & Miller, D. L. Bootstrap-Based Improvements for Inference with Clustered Errors. Review of Economics and Statistics 90 (3), 414–427 (2008). URL https://direct.mit.edu/rest/article/90/3/414-427/57731. https://doi.org/10.1162/rest.90.3.414.

[101] Croft, D. et al. Reactome: A database of reactions, pathways and biological processes. Nucleic acids research 39 (Database issue), D691–7 (2011).

[102] Meyers, R. M. et al. Computational correction of copy-number effect improves specificity of CRISPR-Cas9 essentiality screens in cancer cells. Nat Genet 49 (12), 1779–1784 (2017). URL https://www.ncbi.nlm.nih.gov/pmc/articles/PMC5709193/. https://doi.org/10.1038/ng.3984.

[103] McFarland, J. M. et al. Improved estimation of cancer dependencies from large-scale RNAi screens using model-based normalization and data integration. Nat Commun 9 (1), 4610 (2018). URL https://www.nature.com/articles/s41467-018-06916-5. https://doi.org/10.1038/s41467-018-06916-5.

[104] Korotkevich, G., Sukhov, V. & Sergushichev, A. Fast gene set enrichment analysis. bioRxiv 060012 (2019). URL https://www.biorxiv.org/content/10.1101/060012v2. https://doi.org/10.1101/060012.

[105] Merico, D., Isserlin, R., Stueker, O., Emili, A. & Bader, G. D. Enrichment Map: A Network-Based Method for Gene-Set Enrichment Visualization and Interpretation. PloS one 5 (11), e13984 (2010).

[106] Ba-alawi, W. et al. Bimodal gene expression in cancer patients provides interpretable biomarkers for drug sensitivity. Cancer Research canres.2395.2021 (2022). URL https://doi.org/10.1158/0008-5472.CAN-21-2395. https://doi.org/10.1158/0008-5472.CAN-21-2395.

[107] Bairoch, A. The Cellosaurus, a Cell-Line Knowledge Resource. J Biomol Tech 29 (2), 25–38 (2018). URL https://www.ncbi.nlm.nih.gov/pmc/articles/PMC5945021/. https://doi.org/10.7171/jbt.18-2902-002.

[108] Kundra, R. et al. OncoTree: A Cancer Classification System for Precision Oncology. JCO Clinical Cancer Informatics (2021). URL https://ascopubs.org/doi/pdf/10.1200/CCI.20.00108. https://doi.org/10.1200/CCI.20.00108.

