## Supplementary Figures and Table Captions for "Meta-analysis of preclinical pharmacogenomic studies to discover robust and translatable biomarkers of drug response"

#### Supplementary Tables

**Supplementary Table 1**: All biomarkers (drug-gene-tissue triplets) which had a significant association in at least 1 dataset. The table includes the pearson correlation estimate and p value after meta-analysis.

**Supplementary Table 2**: A table of all genes used in the analysis of Drug Cluster 1 and Cluster 2 associated genesets for Lung Cancer Cell Line Drug Response, with associated cluster assignment. Assignment of a gene to a cluster implies that it predicts sensitivity to drugs in that cluster.

**Supplementary Table 3**: Combined Table of Non-Immunotherapy Datasets identified from the combined CTRDB and SELECT (Lee et al.) resources. Annotation columns represent annotations derived from the CTRDB or Supplementary Information of the Lee et al. manuscript.

**Supplementary Table 4**: The results of the Anderson Darling Test comparing gene expression distribution in cell lines and patient tumours (TCGA).

**Supplementary Table 5**: The results of a permutation test on the concordance index between expression of preclinical biomarkers of Erlotinib response and progression free survival among patients in the Battle1 trial.

**Supplementary Table 6**: The results of Wilcoxon rank-sum test looking at differences in expression of preclinical biomarkers of Lapatinib response among patients in the CHER-LOB trial.

**Supplementary Table 7**: Distribution of diseases among lymphoid cancer cell lines in the datasets considered in this study. See additional files for table.

### Supplementary Figures

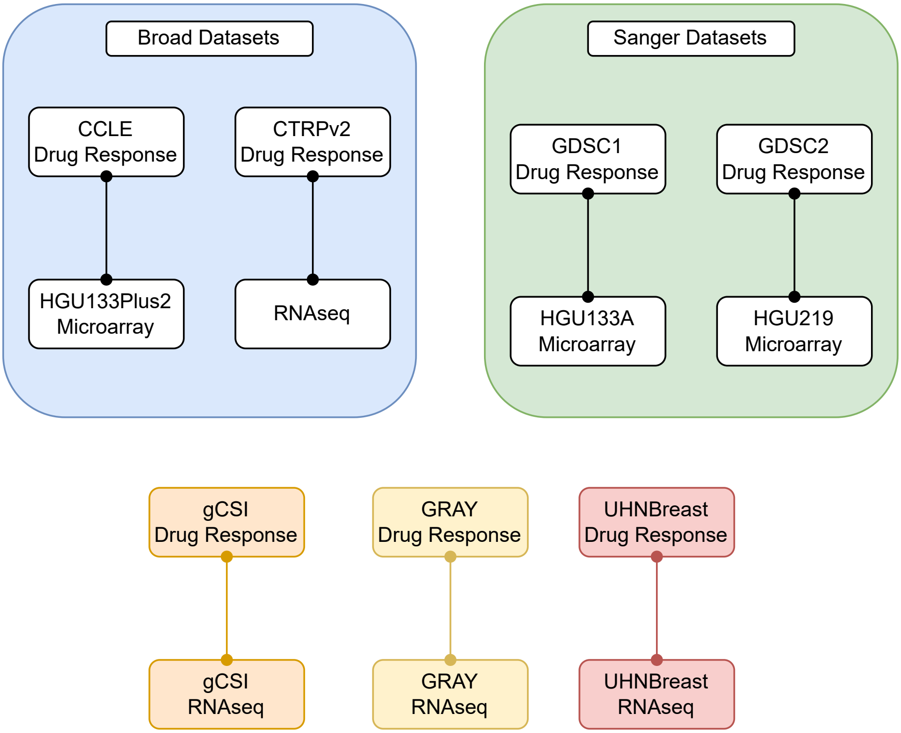


**Supplementary Figure 1**: Diagram displaying the matching between drug response and molecular data, highlighting how data from the Broad and Sanger institutes was matched to create independent datasets.

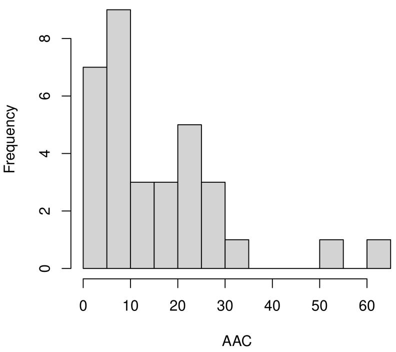


(a)

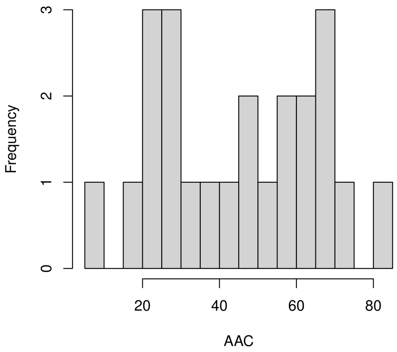


(b)

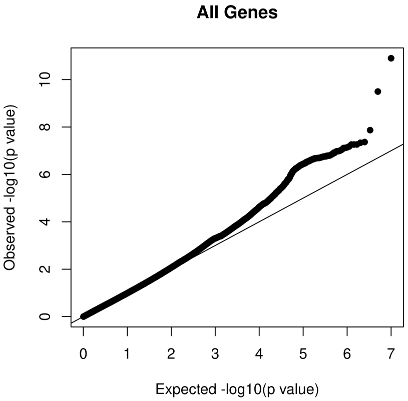


(c)

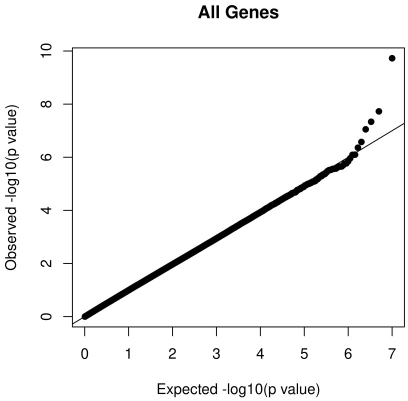


(d)

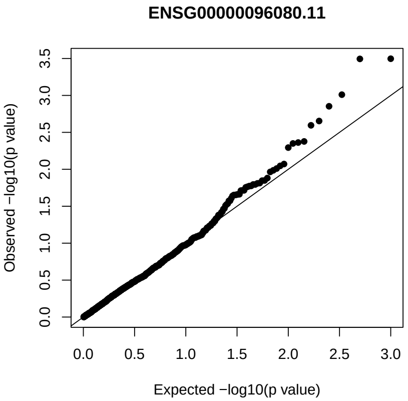


(e)

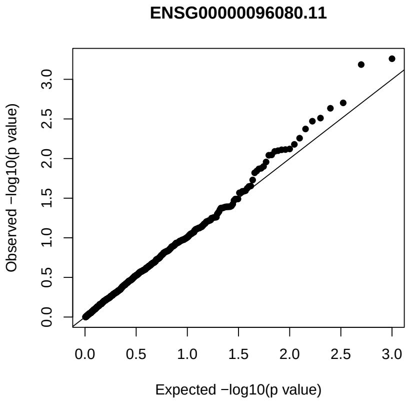


(f)

**Supplementary Figure 2**: Caption on next page.

**Supplementary Figure 2**: **a), b)** The drug response distribution (measured as AAC) for Lapatinib and Paclitaxel respectively among breast cancer cell lines in CTRPv2; **c), d)** Q-Q plots for the expected vs observed distributions of p-values when sampling from a permutation null for associations between gene-expression and drug response tested using the standard Pearson t-test for Lapatinib and Paclitaxel respectively. Results from 10000 permutations of drug response tested against 100 genes shown; **d), e)** Q-Q plots for a specific gene expression association, a subset of the results shown in **c), d)**.

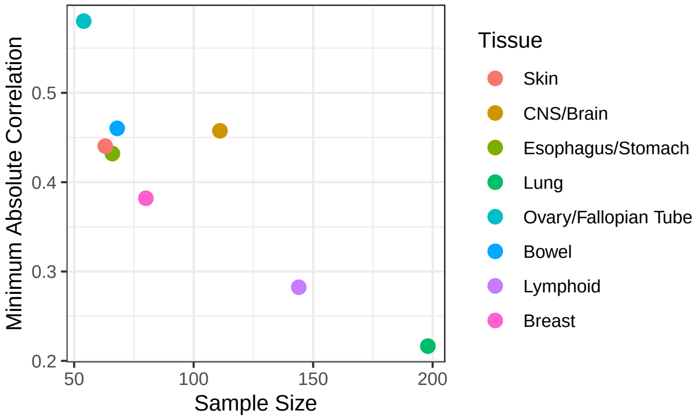


(a)

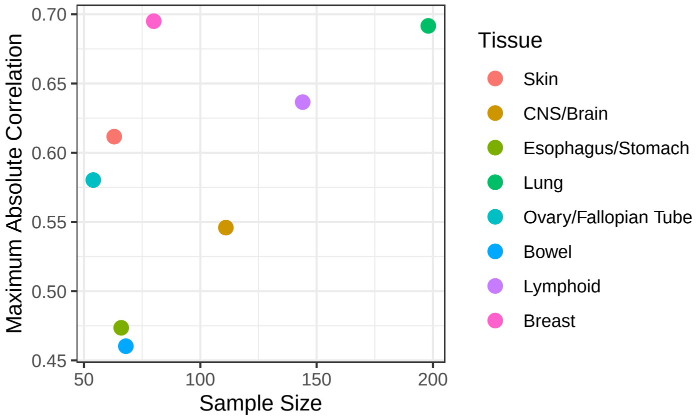


(b)

**Supplementary Figure 3**: **a)** The minimum effect size observed among all significant biomarkers per tissue, as a function of effect size. A significant (Pearson -0.89, p value: 0.004) negative correlation is observed; **b)** the same as a), but showing the maximum effect size. The Pearson correlation of 0.54 is not significant.

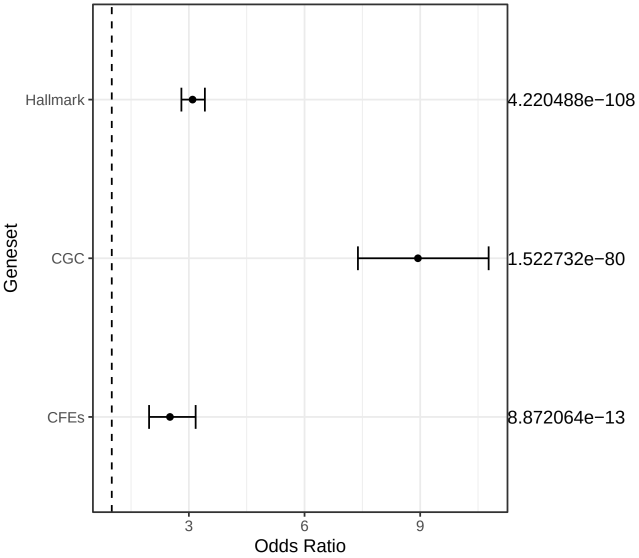


(a)

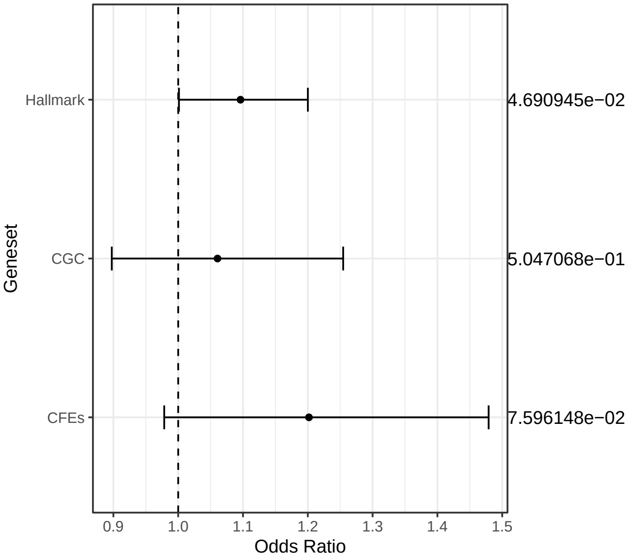


(b)

**Supplementary Figure 4**: Odds ratio's for the enrichment among the Hallmark Genes (mSigDB), Cancer Gene Census (CGC) and Cancer Functional Event genesets of **a)** significant biomarkers of response after meta-analysis, and **b)** proportion of biomarkers passing meta-analysis after being identified as significant in a single dataset.

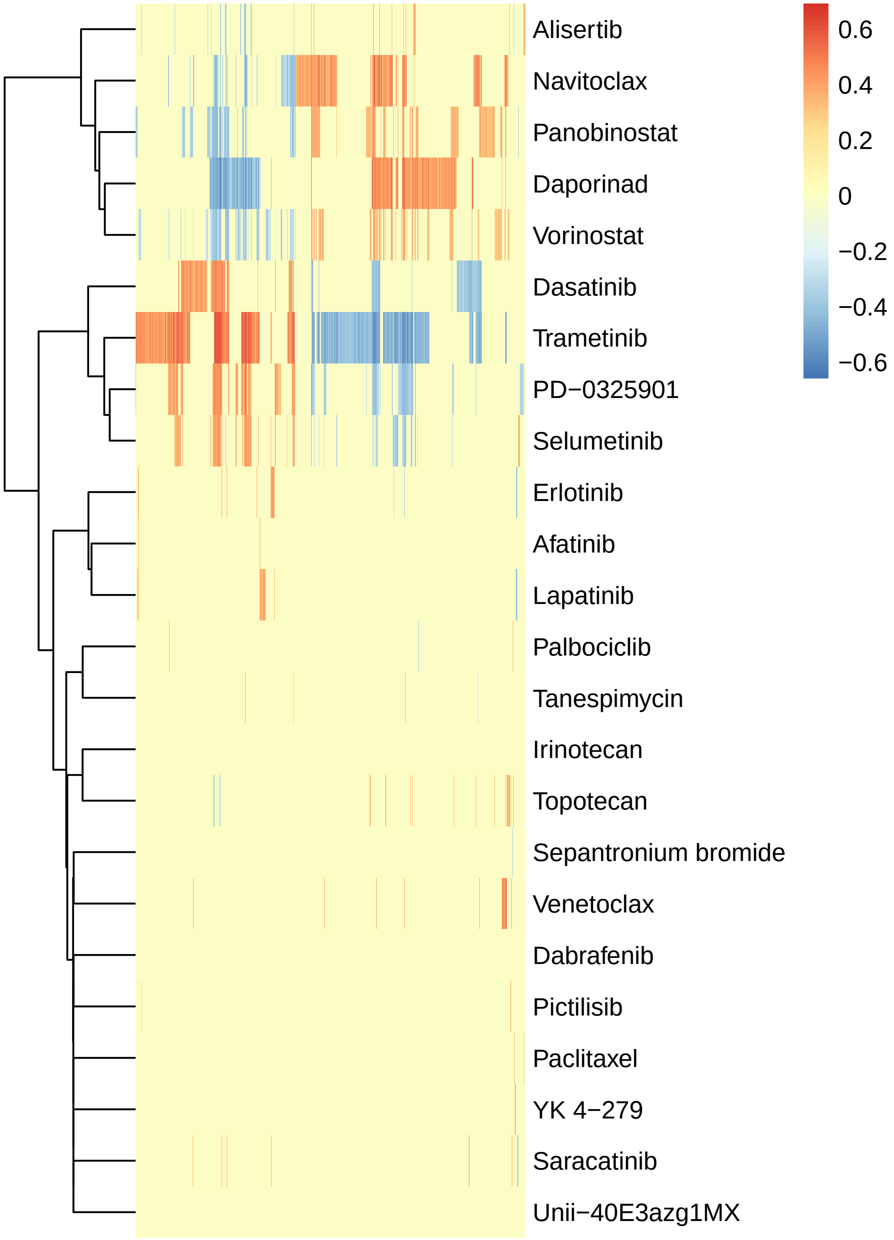


**Supplementary Figure 5**: Heatmap showing the clustering of drugs according to their associations with gene expression after meta-analysis in lung tissue. Non-significant associations are set to 0. Wald D2 clustering on correlation distance is displayed for the rows.

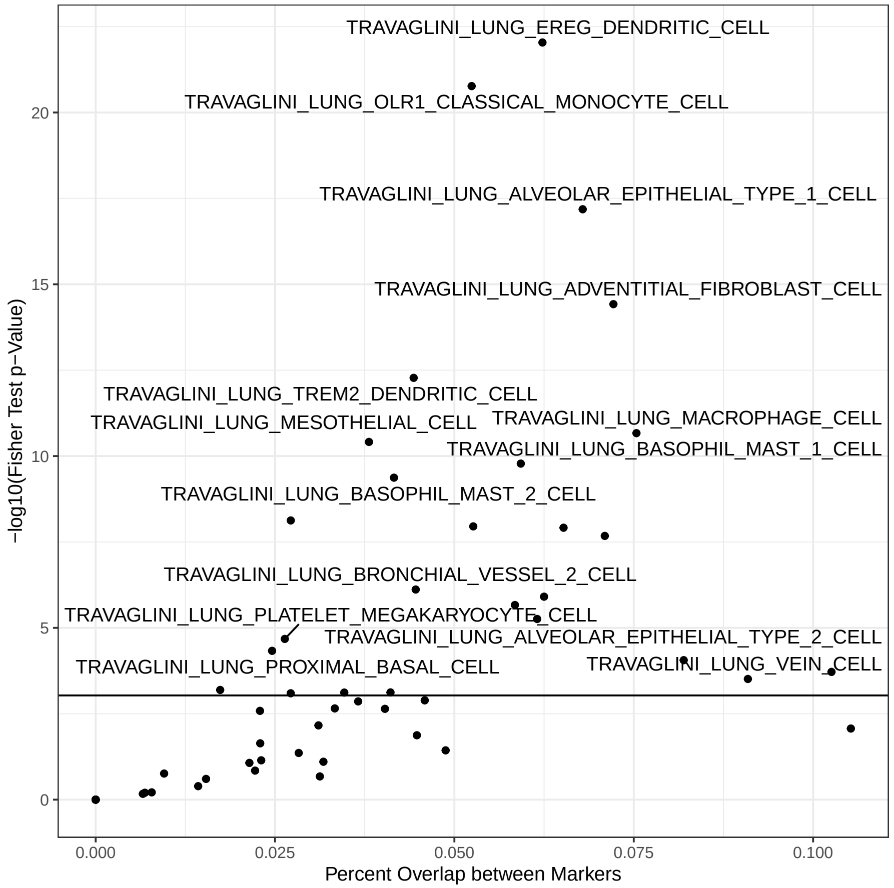


**Supplementary Figure 6**: Enrichment of markers of Lung cancer cell types as defined by Travaglini et al. among biomarkers of sensitivity to cluster 2 drugs/resistance of cluster 1 drugs. The horizontal line indicates the Bonferroni corrected significance cutoff.

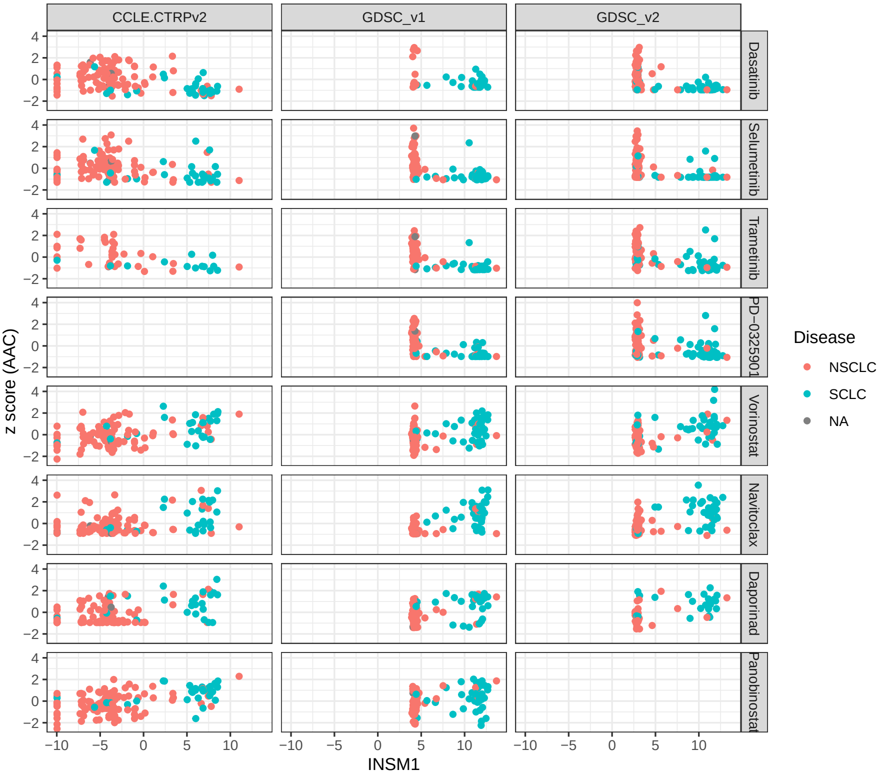


**Supplementary Figure 7**: Standardized drug response plotted against the expression of INSM1 (either in log(TPM+0.001) or log(RMA norm.) scale), colored by the available labels for the histopathologically determined disease from which each cell line originates. Missing panels indicate that the particular drug was not tested in the dataset.

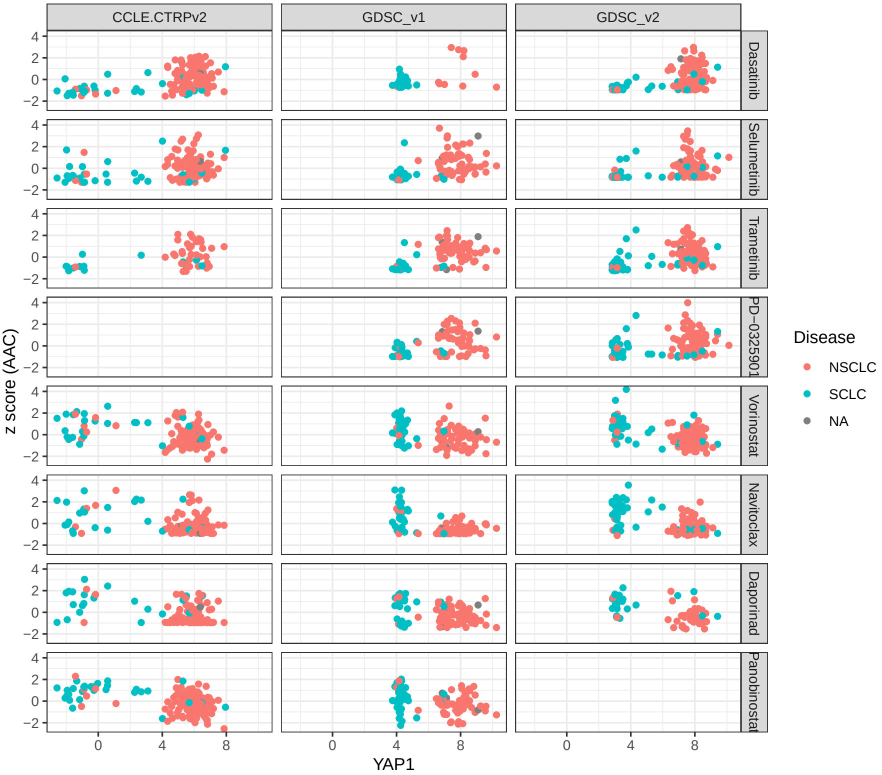


**Supplementary Figure 8**: Standardized drug response plotted against the expression of YAP1 (either in log(TPM+0.001) or log(RMA norm.) scale), colored by the available labels for the histopathologically determined disease from which each cell line originates. Missing panels indicate that the particular drug was not tested in the dataset.

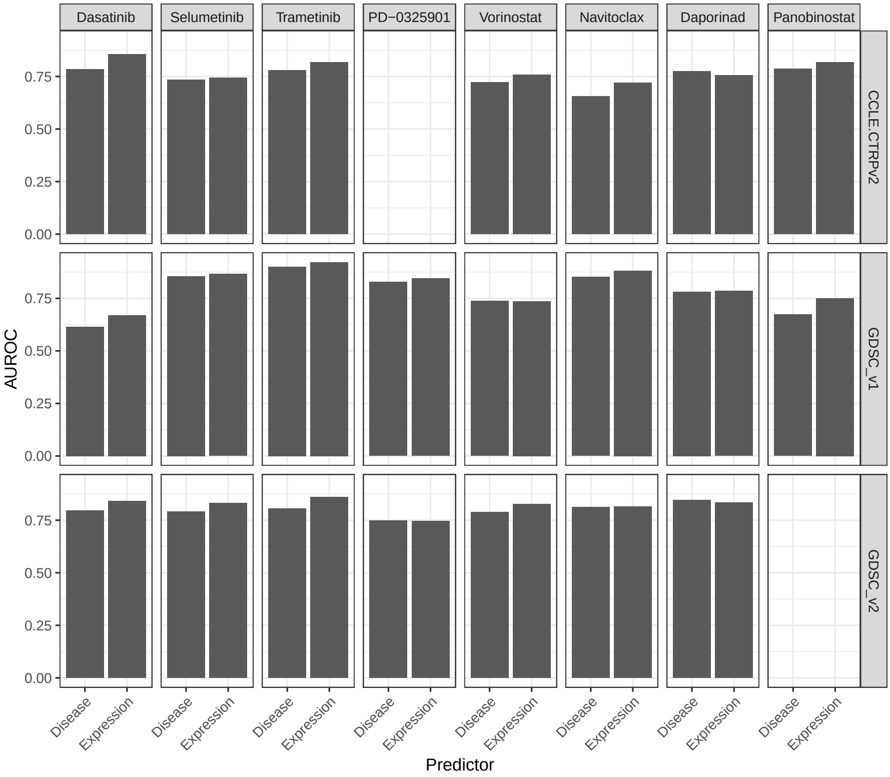


**Supplementary Figure 9**: AUROCs for predicting binary Small Cell Lung Cancer or High INSM1 expression levels from drug AAC values across the 8 drugs and 3 datasets with the majority of drug tested. Class label for cluster 2 drugs is flipped so that AUROC¿0.5 is expected.

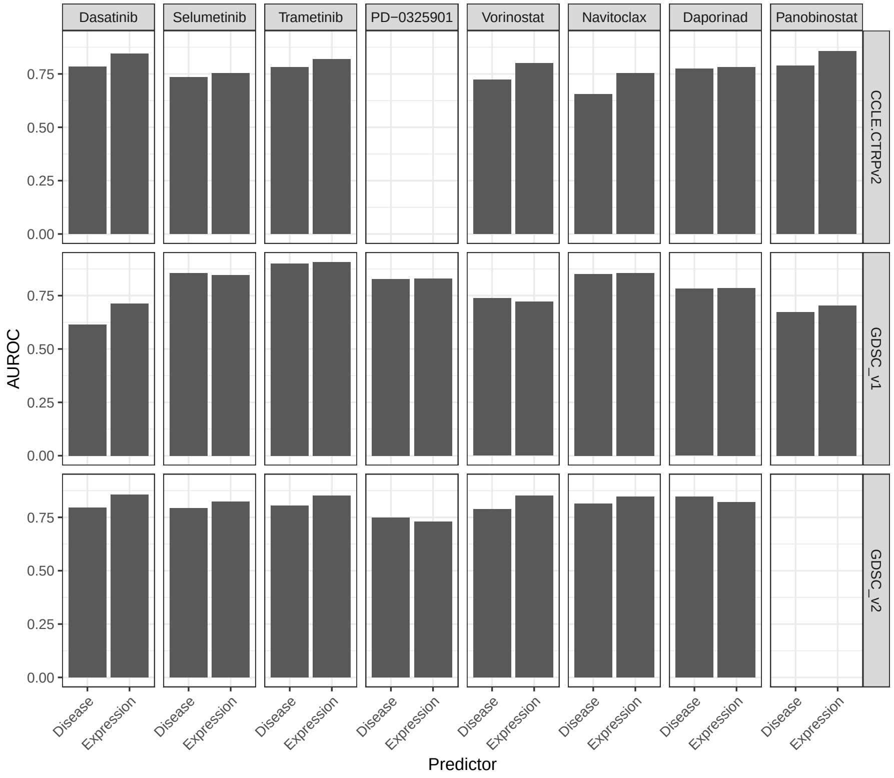


**Supplementary Figure 10**: AUROCs for predicting binary Small Cell Lung Cancer or High YAP1 expression levels from drug AAC values across the 8 drugs and 3 datasets with the majority of drug tested. Class label for cluster 1 drugs is flipped so that AUROC¿0.5 is expected.

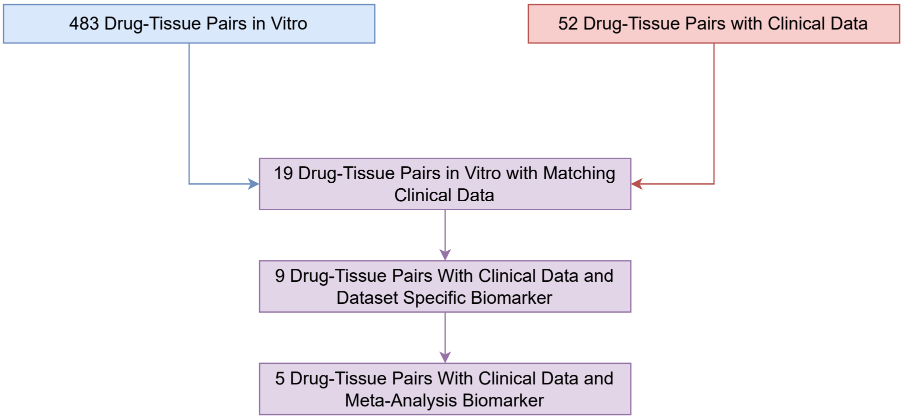


**Supplementary Figure 11**: The results of intersecting the *in vitro* drug-tissue pairs included in the study with available clinical data from the union of datasets in CTR-DB and Lee et al. at different stages of the analysis. Availability of clinical data with drug response and transcriptomics on matching patients remains a major impediment to validating most of the results from our study.

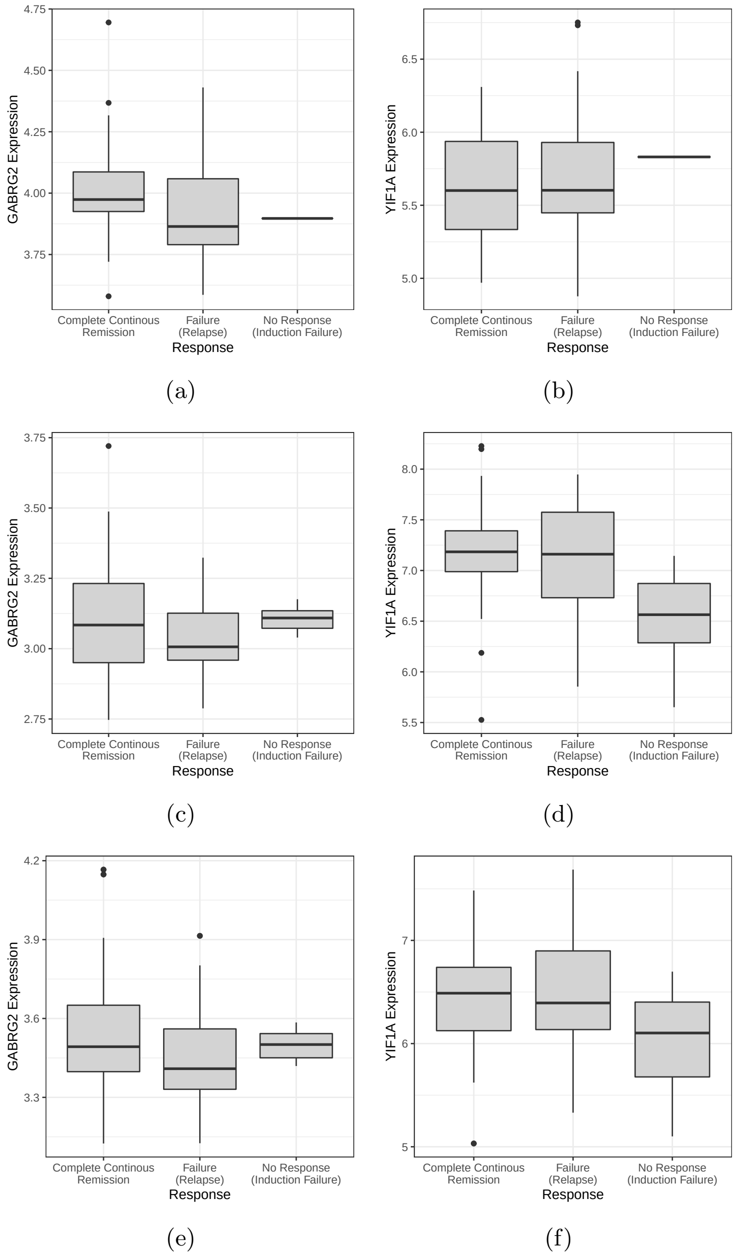


(a) (b) (c) (d) (e) (f)

**Supplementary Figure 12**: **a),b)** The expression (log(RMA) units) of GABRG2 and YIF1A across different response groups in the COG 8704 study; **c),d)** The expression (log(RMA) units) of GABRG2 and YIF1A across different response groups in the COG 9404 study; **e),f)** ComBat corrected expression (log(RMA) units) of GABRG2 and YIF1A across different response groups in COG9404 and COG8704 together.

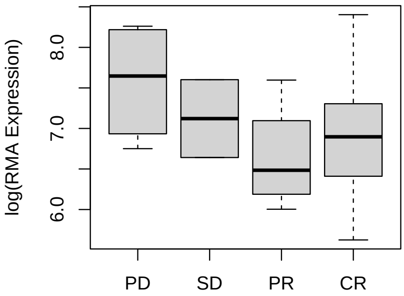


(a)

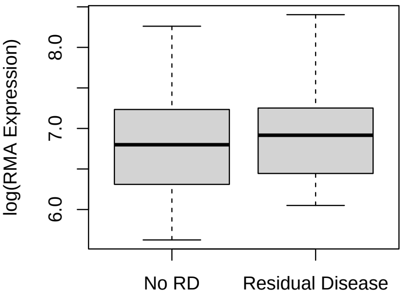


(b)

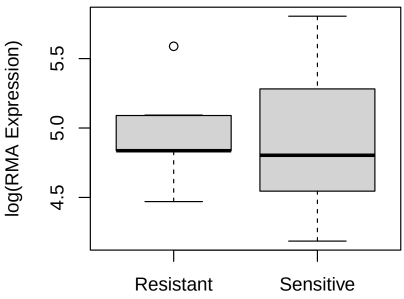


(c)

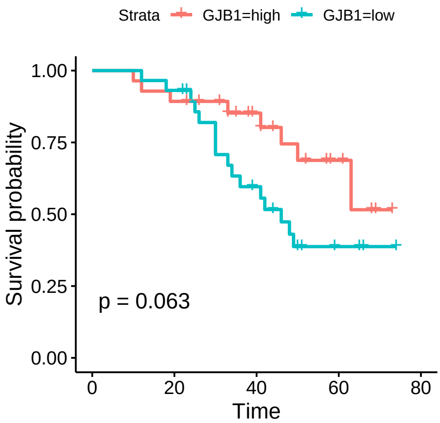


(d)

**Supplementary Figure 13**: Relationship between GJB1 expression and response among the **a)** GSE63885, **b)** GSE14764 and **c)** GSE15622 datasets, with no significant associations; **d)** the association between GJB1 expression and survival within the GSE31245 dataset.

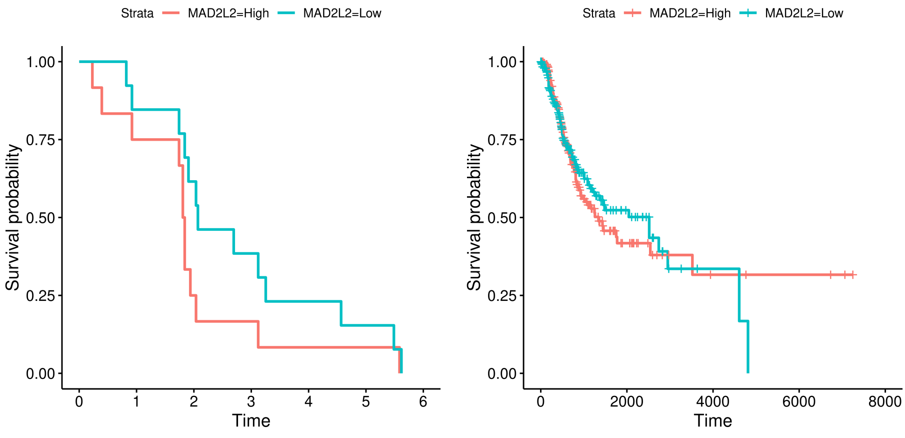


(a) (b)

**Supplementary Figure 14**: The Kaplan Meier curves highlighting the association between MAD2L2 expression and survival among **a)** BATTLE1 trial patients (unadjusted p value: 0.0078) and **b)** the LUAD TCGA cohort (Not Significant).

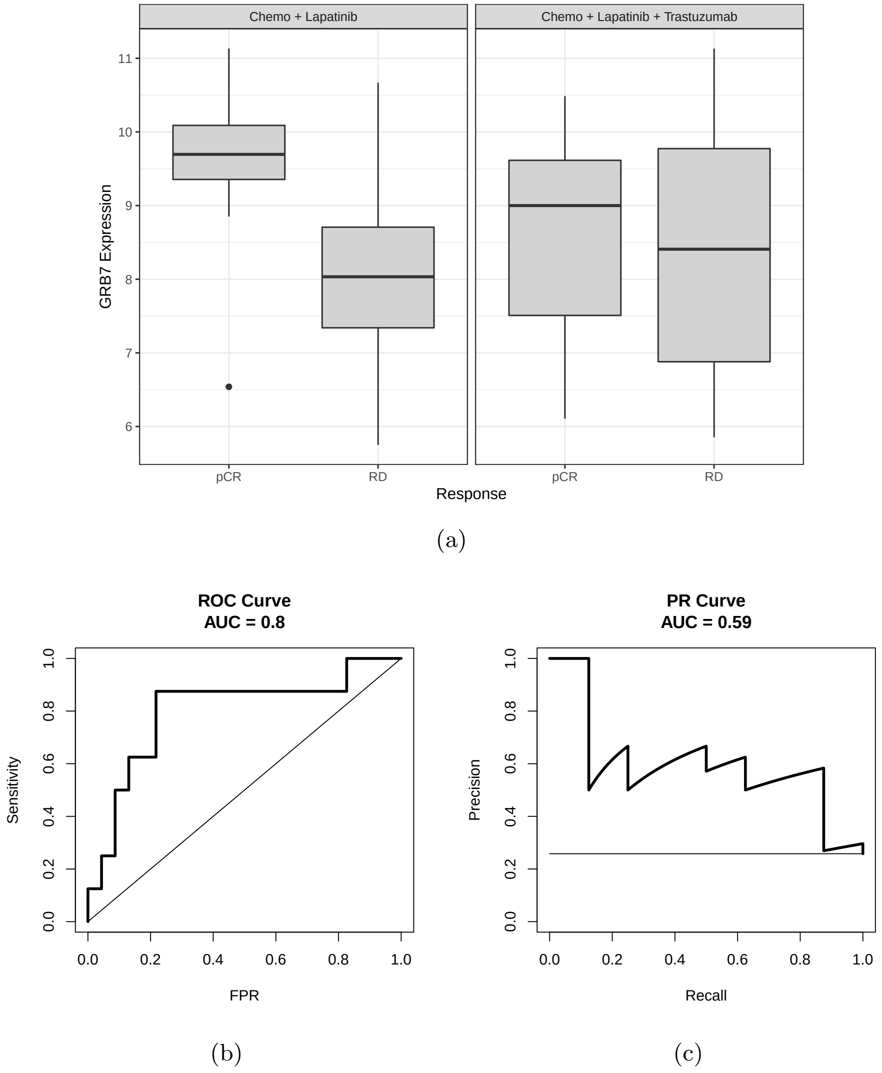


**Supplementary Figure 15**: **a)** Expression of GRB7 among responding and non-responding patients in the CHER-LOB study, split by treatment arm for those arms containing Lapatinib; **b),d)** The ROC and PR curves for GRB7 as a predictor of pCR among the Chemo + Lapatinib arm of the CHER-LOB study;

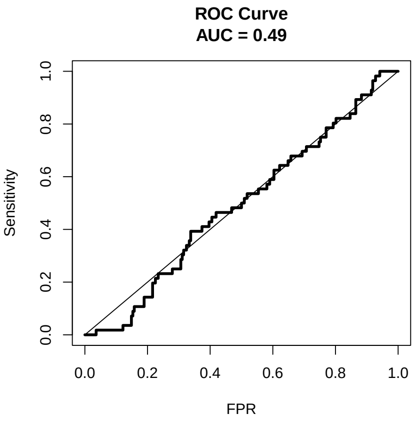


(a)

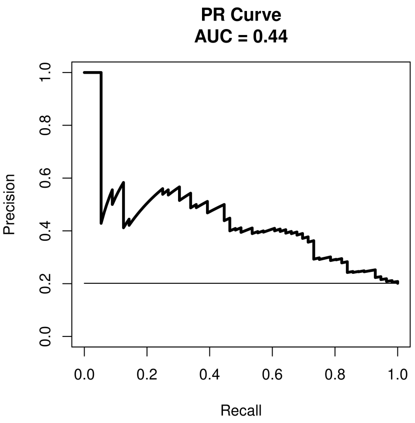


(b)

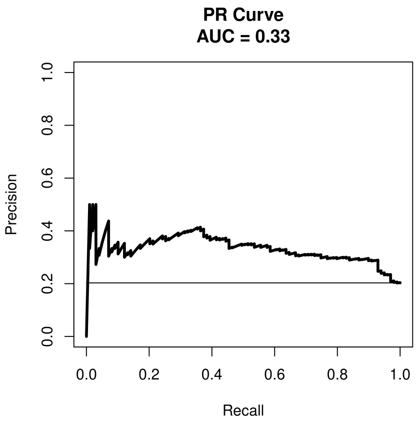


(c)

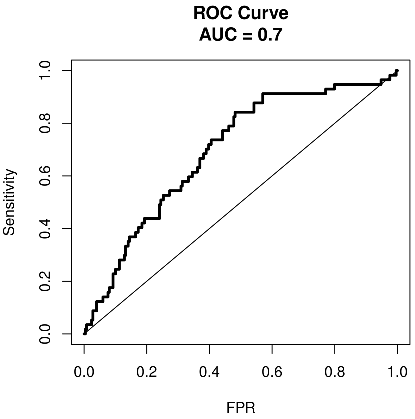


(d)

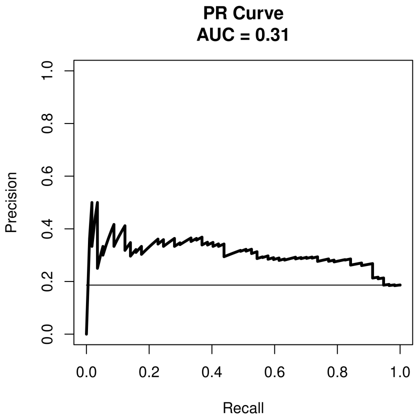


(e)

**Supplementary Figure 16**: Caption on next page

**Supplementary Figure 16**: **a)** ROC curve for EIF5A expression as a predictor of pCR in the MAQCII dataset; **b),c)** PR curves for ODC1 expression as a predictor of pCR in the MAQCII and Hatzis datasets respectively;**d),e)** ROC and PR curves (respectively) for ODC1 expression as a predictor of pCR in the Hatzis discovery sub-cohort;

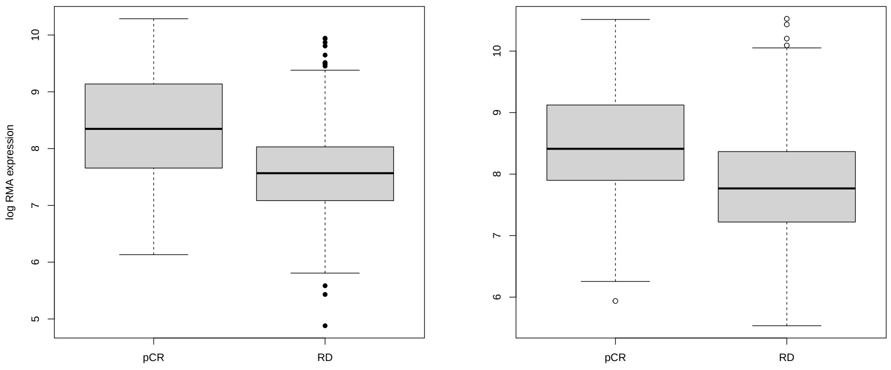


(a) (b)

**Supplementary Figure 17**: **a),b)** Expression of ODC1 in the MAQCII and Hatzis datasets split between responders and non-responders.

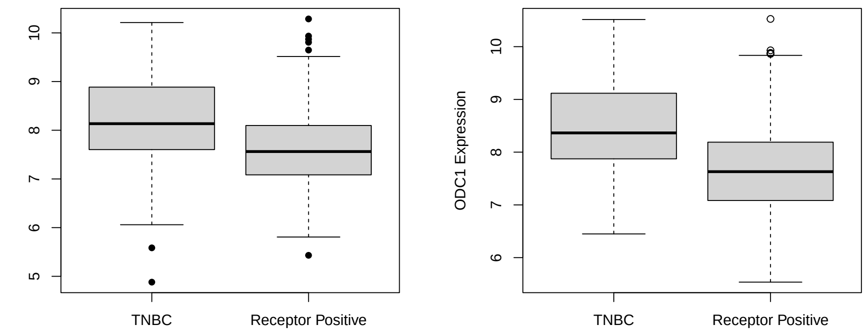


(a) (b)

**Supplementary Figure 18**: **a), b)** Expression of ODC1 in the MAQCII and Hatzis datasets split between TNBC and receptor positive samples.

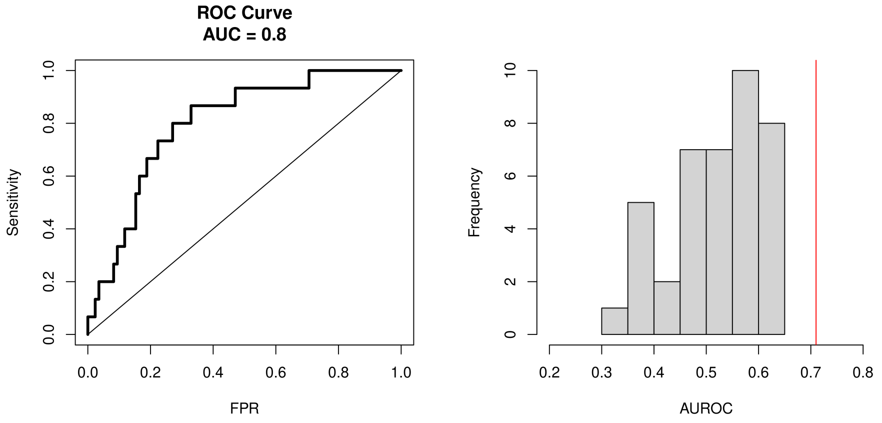


(a) (b)

**Supplementary Figure 19**: **a)** ROC curve for performance of ODC1 expression as a predictor of pCR for patients in the validation subset of the data; **b)** Distribution of predictors of pCR trained and evaluated on ER- patients in the MAQCII study compared to the performance of ODC1 as a predictor for ER- patients.
